# Quiescent Cells Exhibit a Senescence-Like Secretory Phenotype

**DOI:** 10.64898/2026.11.15.766855

**Authors:** Azucena Rocha, Radha Kalekar, John Sedivy, Nicola Neretti

## Abstract

Cellular quiescence is traditionally considered a reversible state of cell cycle arrest, distinct from senescence, which is associated with a characteristic senescence-associated secretory phenotype (SASP). However, whether quiescent cells exhibit a comparable secretory program remains poorly understood. Here, we identify and characterize a Quiescence-Associated Secretory Phenotype (QASP) by integrating single-cell and bulk RNA sequencing datasets from human fibroblasts and preadipocytes. We find that quiescent cells exhibit increased expression of genes encoding inflammatory mediators and extracellular matrix-associated proteins, with substantial overlap between the QASP and SASP. This transcriptional program is highly heterogeneous, varying across cell types, quiescence induction methods, and environmental oxygen levels. In particular, contact inhibition induces a stronger inflammatory signature than serum withdrawal in several cellular contexts. Multiplexed cytokine profiling demonstrated active secretion of inflammatory mediators, including IL-1*β*, IL-6, IL-15, and CCL2, in quiescent fibroblasts, with secretion levels varying by induction method and generally higher following contact inhibition than serum withdrawal. Analysis of single-cell transcriptomic data from human lung tissue further reveals increased QASP-associated gene expression in idiopathic pulmonary fibrosis (IPF), particularly in myofibroblasts, linking this phenotype to inflammatory signaling and extracellular matrix remodeling. Together, these findings identify the QASP as a context-dependent secretory program that shares features with the SASP and may contribute to inflammatory signaling and extracellular matrix remodeling in both physiological and pathological conditions.

## Introduction

Cellular quiescence is a reversible state of growth and cell-cycle arrest that cells undergo in response to diverse environmental stimuli, such as contact inhibition (CI), nutrient deprivation, mitogen withdrawal, oxidative stress, and DNA damage. Quiescent cells exit the cell cycle in the G0 phase but remain metabolically active and can re-enter the cell cycle as needed [1]. They are critical for maintaining tissue homeostasis and are found in many tissues throughout the body, particularly in stem cell, fibroblast, and immune cell lineages [2, 3]. Quiescent cells exhibit distinct expression patterns compared to proliferating cells, especially for pathways related to cell cycle progression, tumor suppression, and extracellular matrix (ECM) remodeling [4, 5, 6]. Several experimental models have been developed to induce quiescence *in vitro*. CI [7] is induced by allowing cells to reach confluence, which stalls growth due to reduced access to growth-promoting mitogens. Another commonly used method is serum withdrawal (SW) [8], which deprives the cells of essential growth factors, halting cell cycle progression. While quiescence is typically associated with reduced cellular activity, evidence suggests that quiescent cells actively engage in tissue maintenance and response to stress by secreting factors that influence the local tissue environment [4]. However, the extent and nature of this secretory activity remain poorly defined.

Cellular senescence is another form of cell cycle arrest, characterized by irreversible exit from the cell cycle in response to cellular stressors resulting in DNA damage-induced senescence (DDIS), oncogene-induced senescence (OIS) or telomere shortening, known as replicative senescence (RS) [9, 10, 11]. Unlike cellular quiescence, cellular senescence is irreversible. However, recent studies have questioned whether quiescence and senescence are binary, distinct cellular states or whether they exist on a continuum of cell-cycle withdrawal [12].

The secretome refers to the collection of proteins secreted by cells into the extracellular space, which plays a central role in cell-cell communication, tissue remodeling, and immune response. Approximately 36% of human protein-coding genes have predicted secreted transcripts, and 1891 genes are predicted to have at least one secreted protein product [13]. These secreted proteins, including cytokines, growth factors, and other signaling molecules, play crucial roles in many physiological and pathological processes.

A key characteristic of senescent cells is an inflammatory signature known as the Senescence-Associated Secretory Phenotype (SASP) [14]. The SASP plays a dual role in tissue repair and chronic inflammation. In contrast, the secretory activity of quiescent cells has not been extensively characterized. While several studies have demonstrated differences in gene expression between proliferating and quiescent cells, and that quiescent fibroblasts mount a more robust inflammatory response compared to their proliferating counterparts [15], there is limited data describing the secretory profile in these cells.

In this study, we used a combination of single-cell transcriptomics (scRNA-Seq) on human lung fibroblasts, and bulk RNA sequencing (bulk RNA-Seq) to identify a putative secretory signature in quiescent cells. From a previously published list of 1891 genes predicted to encode the secretome [13], we selected those that were transcriptionally upregulated in quiescent compared to proliferating cells. We refer to this subset of genes as the transcriptional Quiescent-Associated Secretory Phenotype (QASP), drawing an analogy to the SASP, where genes are similarly transcriptionally upregulated during senescence.

We complemented our transcriptomic data with a multiplex cytokine assay to directly measure cytokine secretion. We observed increased secretion of several key cytokines, including *IL6*, *IL1B*, *IL15*, and *CCL2*, in the media of CI-induced quiescent fibroblasts. This confirms that QASP is not just a transcriptional phenomenon but reflects a functional inflammatory secretory phenotype—supporting the existence of a true Quiescence-Associated Secretory Phenotype (QASP).

We combined in-house generated scRNA-Seq and bulk RNA-Seq datasets of quiescent and senescent human fibroblasts and preadipocytes cultured under different oxygen levels with publicly available data on senescent fibroblasts, and analyzed the expression patterns of QASP genes. Additionally, we compared these patterns with those observed in senescent human fibroblasts at different time points. Finally, we validated our findings by identifying the QASP in publicly available scRNA-Seq data from pulmonary fibrotic tissue, providing insights into the role of quiescence in tissue pathology. This comprehensive analysis enabled a unified examination of inflammation in quiescence, offering novel insights into its implications in disease pathology and aging.

## Results

### Single-cell RNA profiling reveals an inflammatory transcriptional profile in quiescent LF1 fibroblasts

Our previously published study on cell-to-cell transcriptional heterogeneity in different forms of senescence (RS, OIS, and DDIS) used proliferating cells as controls [16]. To compare senescence heterogeneity with a different form of cell cycle arrest, we performed scRNA-seq on LF1 lung fibroblasts induced into quiescence with a combination of CI and SW (Fig. 1 A). As expected, quiescent cells clustered separately from proliferating and all forms of cellular senescence (Fig. 1 B). These cells exhibited downregulation of the proliferation marker *MKI67*, upregulation of the quiescent associated CDK inhibitors *CDKN1B* and *CDKN1C* [17], and lower expression levels of the senescence markers *CDKN1A* and *CDKN2A* with respect to senescent cells, while maintaining expression levels comparable to those of proliferating cells (Fig. 1 C). Unexpectedly, many quiescent cells showed significantly higher expression levels of the pro-inflammatory cytokines *IL6* and *IL1B* with respect to all forms of senescence (Fig. 1 C and D). To ensure these cells did not represent a subpopulation of senescent cells within the quiescent cell culture, we excluded cells expressing *CDKN2A* and *CDKN1A* from the quiescent sample. Even after this filtration, the levels of inflammatory cytokines remained significantly higher than those in senescent cells (Supplemental Fig. 1), indicating that the observed pro-inflammatory phenotype is an intrinsic characteristic of quiescent cells, rather than a contamination by senescent cells.

**Figure 1.**
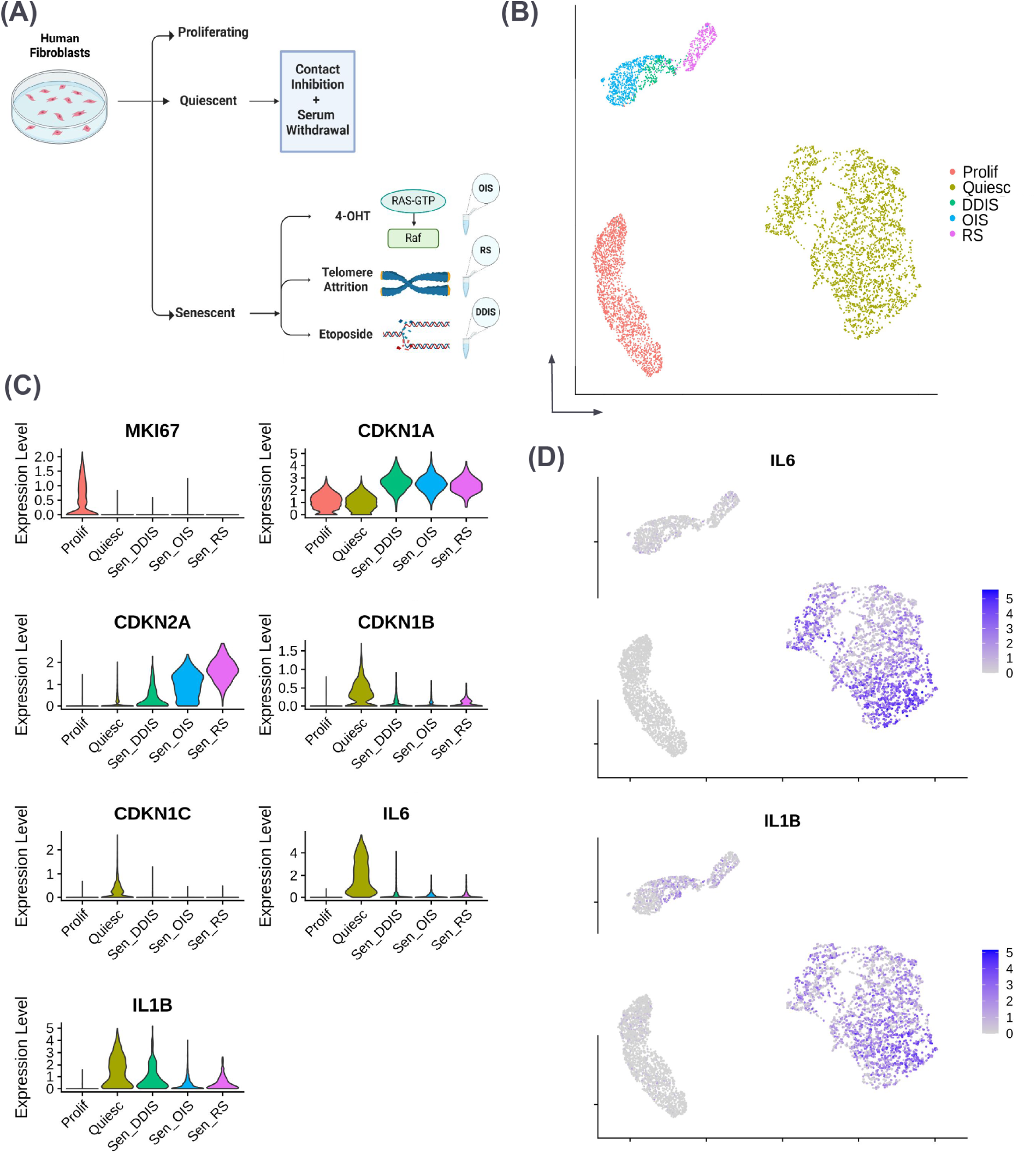
Single-cell RNA sequencing analysis of quiescent and senescent samples. (A) Schematic representation of the experimental design, illustrating the generation of single-cell RNA sequencing libraries from various conditions (proliferating, quiescent, and senescent). Senescent cells were induced via replicative exhaustion (RS), etoposide treatment (DDIS), or oncogenic activation (OIS). Single-cell libraries were generated using 10x Chromium technology. (B) Uniform Manifold Approximation and Projection (UMAP) plots displaying all datasets generated for this study, showing distinct clustering of proliferating, quiescent, and senescent cells. (C) Violin plots showing lack of the proliferation marker *MKI67* in quiescent and senescent cells, and expression of inflammatory genes *IL6* and *IL1B* in proliferating, quiescent, and senescent cells. (D) Feature plots showing increased expression of *IL6* (top) and *IL1B* (bottom) in the quiescent cell cluster.

To further explore this unexpected finding, we constructed a comprehensive list of significant differentially expressed genes from our quiescent single-cell and bulk RNA-Seq data from five different cell lines: BJ fibroblasts, IMR90 fibroblasts, LF1 fibroblasts, WI38 fibroblasts, and primary preadipocytes compared to proliferating cells (Table 1). Quiescence was induced via CI, SW, or both. Quiescent cells were maintained at either atmospheric (20%) or physiological (2.5%) O_2_ levels. Significance was determined as false discovery rate (FDR) *<* 0.05 and log fold change (log_2_FC) *>* 1. From this pool of significantly expressed genes, we extracted those transcriptionally upregulated from the list of 1891 genes in the human secretome [13]. This subset of genes represents the putative QASP. We then performed differential gene expression analysis on our single-cell data, comparing quiescent or senescent cells (DDIS, OIS, or RS) to proliferating controls and extracted the genes present in our QASP list. A substantial proportion of quiescent cells exhibited upregulated expression of QASP genes, displaying a distinct transcriptional profile despite some overlap with senescent cells. Senescent cells clustered together, demonstrating a more similar expression pattern to each other than to quiescent cells (Supplemental Fig. 2B).

**Table 1:** Datasets, Cell Line, Induction Method, and Oxygen Levels used to generate the QASP and the SASP.

| **Category** | **Dataset Type** | **Cell Line** | **Induction Method** | **Oxygen Level** |
|---|---|---|---|---|
| SASP | scRNA-Seq | LF1 | DDIS | low ox |
|  | scRNA-Seq | LF1 | OIS | low ox |
|  | scRNA-Seq | LF1 | RS | low ox |
|  | Bulk RNA-Seq | BJ | RS | high ox |
|  | Bulk RNA-Seq | IMR90 | RS | high ox |
|  | Bulk RNA-Seq | LF1 | Early RS | low ox |
|  | Bulk RNA-Seq | LF1 | Late RS | low ox |
| QASP | scRNA-Seq | LF1 | CI + SW | low ox |
|  | Bulk RNA-Seq | BJ | CI | high ox |
|  | Bulk RNA-Seq | BJ | SW | high ox |
|  | Bulk RNA-Seq | Preadipocytes | CI | high ox |
|  | Bulk RNA-Seq | Preadipocytes | SW | high ox |
|  | Bulk RNA-Seq | IMR90 | CI | high ox |
|  | Bulk RNA-Seq | IMR90 | SW | high ox |
|  | Bulk RNA-Seq | WI38 | CI | high ox |
|  | Bulk RNA-Seq | BJ | CI | low ox |
|  | Bulk RNA-Seq | BJ | SW | low ox |
|  | Bulk RNA-Seq | LF1 | SW | low ox |

Our analysis of scRNA-Seq data demonstrates that quiescent LF1 fibroblasts exhibit an unexpected expression of pro-inflammatory cytokines *IL6* and *IL1B*. Using this data in combination with bulk RNA-Seq from quiescent BJ and IMR90 fibroblasts and preadipocytes, we mapped the significantly upregulated genes also found in the secretome and defined this as the QASP. We found that a large proportion of quiescent cells exhibited upregulated expression of the QASP, displaying a transcriptional profile distinct from senescent cells. These findings suggest that quiescent cells may participate in physiological and inflammatory signaling processes previously unrecognized, potentially redefining our understanding of the role in quiescence in cellular homeostasis and stress responses.

### Quiescent Cells Exhibit Distinct, Induction-Dependent Transcriptional Profiles Across Cell Types

To determine whether the QASP expression in quiescent cells is specific to cell line or induction method, we performed bulk RNA-Seq on three different cell lines: BJ fibroblasts, IMR90 fibroblasts, and primary preadipocytes). Quiescence was induced using either CI or SW, the cells were maintained under atmospheric (*∼* 20%) oxygen levels and all samples were performed in triplicate (Fig. 2A).

**Figure 2.**
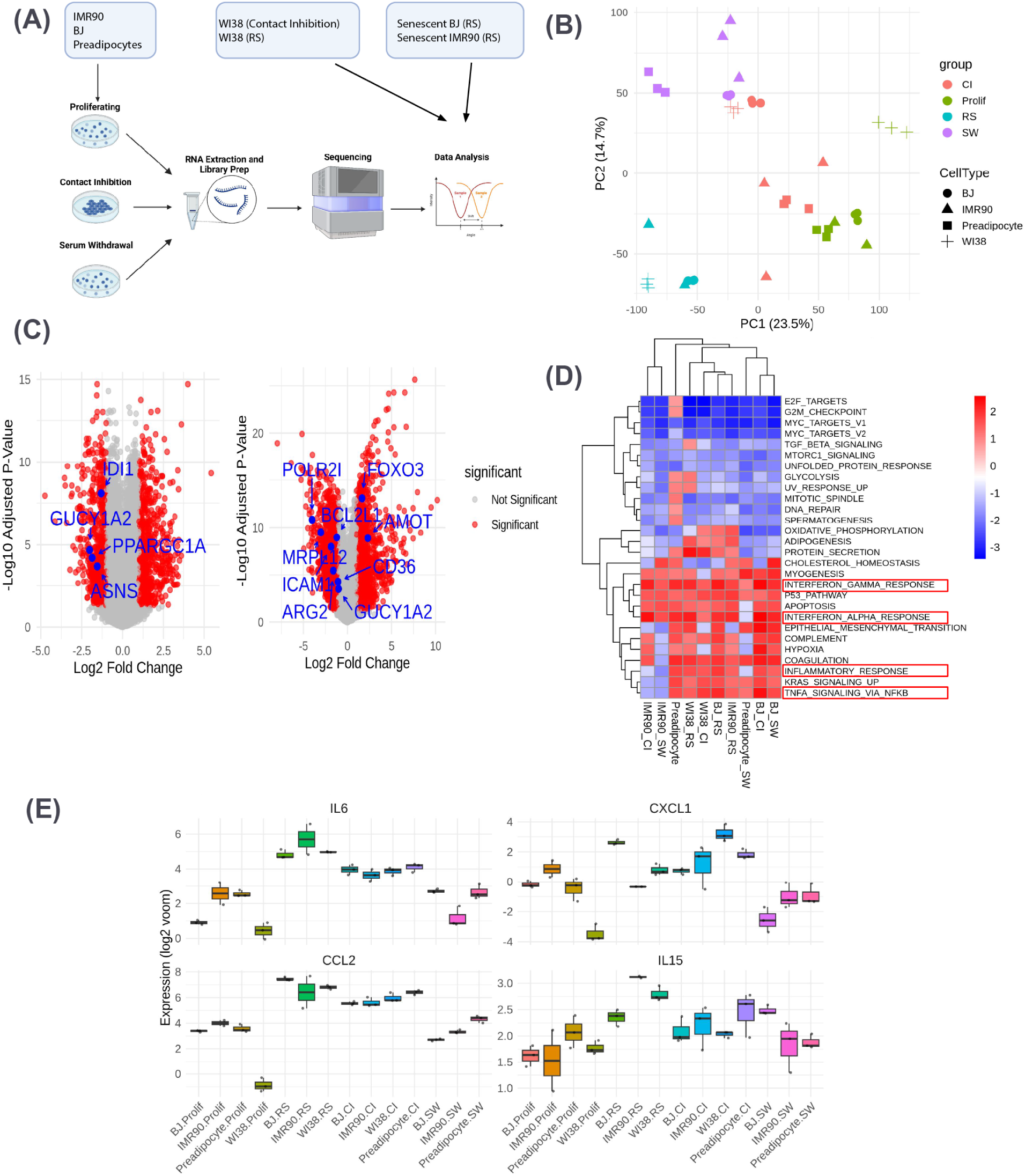
Experimental design and comparative analysis of gene expression across four cell lines, two quiescence induction methods, and replicative senescence. (A) Schematic representation of the bulk RNA-seq experimental design, illustrating the four cell lines (BJ, IMR90, WI38, and preadipocytes) and four conditions (proliferation, contact inhibition, serum withdrawal, and replicative senescence). (B) PCA plot highlighting the separation of samples by cell type and condition after batch correction. BJ cells are depicted as circles, IMR90 cells as triangles, WI38 cells as plus signs, and preadipocytes as squares. CI is shown in red, Prolif in green, RS in blue, and SW in purple. (C) Volcano plots illustrating upregulated and downregulated genes in each comparison. Highlighted genes are known to change in CI or SW (left) or in quiescence versus senescence (right). (D) GSEA heatmap showing differentially upregulated and downregulated pathways across cell lines and conditions for significantly differentially expressed genes (FDR *<* 0.05 and log_2_ FC *>* 1) compared with proliferating cells. Inflammatory pathways are highlighted. (E) Box plots showing expression (log_2_ normalized) of inflammatory genes *IL6*, *CXCL1*, *CCL2*, and *IL15* in CI, SW, proliferating, and senescent cells across LF1 and IMR90 cell lines cultured under low-oxygen conditions.

We incorporated publicly available bulk RNA-Seq data from the Gene Expression Omnibus (GEO) repository (accession number GSE63577) into our analysis, providing transcriptomic profiles of senescent human fibroblasts generated from IMR90 and BJ cells that underwent replicative exhaustion (RS) maintained under atmospheric (*∼* 20%) oxygen levels[18]. We also incorporated publicly available bulk RNA-Seq data from GEO repository (accession number GSE175533), providing transcriptomic profiles of quiescent (CI) and senescent (RS) WI38 fibroblasts maintained under atmospheric (*∼* 20%) oxygen levels [19]. These data were processed in parallel with our own using the same normalization and differential expression pipelines to ensure consistency, and batch correction was applied to the combined dataset. Principal Component Analysis (PCA) revealed clustering by condition, with SW-induced samples clustering in the top left, RS samples in the bottom left, CI-induced samples in the middle, and proliferating samples on the right (Fig. 2B). The two quiescence induction methods clustered closer together along PC1 than they did to proliferating cells, and were separated from senescent cells along PC2. Non-batch corrected data are presented in Supplemental Fig. 2A.

Differential gene expression analysis was performed for each cell line between quiescence (CI or SW-induced) or senescence compared to proliferating controls, resulting in ten comparisons (three for BJ cells, three for IMR90, two for WI38, and two for preadipocytes). A heatmap of the log_2_FC of significantly differentially expressed genes (adjusted p-value *<* 0.05, absolute log_2_FC *>* 1) highlighted common and unique gene expression patterns among quiescence induced by CI and SW and senescence. While CI and SW-induced cells displayed more similar expression patterns to each other than senescent cells, each quiescence induction method showed unique expression profiles across all cell lines. Notably, preadipocytes showed the most pronounced expression pattern differences between CI and SW (Supplemental Fig. 2B).

To analyze differentially expressed genes between conditions, we generated two volcano plots (Fig.2C) to compare CI with SW-induced cells, and quiescence (combined CI and SW induction vs. senescence (RS), respectively. Analyses were performed on merged data from BJ and IMR90 cells. In the CI vs. SW comparison, genes such as *IDI1*, which is involved in lipid metabolism, and *GUCY1A2*, involved in signal transduction, and genes involved in amino acid metabolism such as *ASNS* and *PPARGCI*, were upregulated in SW, consistent with previous reports [4]. The comparison between quiescence and senescence revealed downregulation of genes such as *POLR2I* and *MRPL12*, involved in protein metabolism and *GUCYIA2*, in quiescent cells. Senescence-associated cell surface marker genes like *ICAM1* and *CD36* showed upregulation in senescence [20]. Quiescence-specific genes such as *FOXO3* and *AMOT* were also upregulated in quiescence [21, 22, 23]. These comparisons confirm that distinct gene expression patterns are associated with different quiescence induction methods and highlight transcriptional differences between quiescence and senescence.

To further explore the biological pathways that are upregulated or downregulated between experimental groups, we performed Gene Set Enrichment Analysis (GSEA) for significantly differentially expressed genes in comparisons between CI vs. proliferating and SW vs. proliferating cells for each of the three cell lines, as well as RS vs. proliferating for BJ and IMR90 cells (Fig.2D). We observed upregulation of inflammatory pathways, including IL6-JAK-STAT signaling, inflammatory response, and interferon *alpha* and *gamma* response pathways in both the RS vs. proliferating comparisons and CI vs. proliferating comparisons for preadipocytes, IMR90, and BJ cells, as well as SW vs. proliferating for BJ cells. This upregulation suggests a shift towards a pro-inflammatory state in these conditions. Additionally, TNF-*alpha* signaling via NF-*κ*B was consistently upregulated in all comparisons involving BJ cells and preadipocytes but was not upregulated in quiescent IMR90 cells, indicating potential cell line-specific differences in the inflammatory response during quiescence. Box plots of specific inflammatory genes, including *IL6, CXCL1, CCL2* and *IL15*, confirmed that the gene expression patterns are heavily influenced by the quiescence induction methods. Most CI samples showed high and similar expression levels, while most SW samples expression levels were lower, although these differences were cell line dependent. Proliferating samples consistently showed lower expression levels than quiescence and senescence, while RS samples exhibited high expression patterns across different cell lines (Fig. 2E).

Finally, we examined the expression of cell cycle inhibitors for each condition. Consistent with our previous findings, we observed upregulation of *CDKN1B* (p27) and *CDKN1C* (p57) in quiescence, particularly in the SW condition (Supplemental Fig. 2C). Additionally, *CDKN2B* (p15) expression was higher in SW, whereas *CDKN2D* (p19) was higher in CI. Both *CDKN1A* (p21) and *CDK2N2A* (p16) exhibited high heterogeneity in expression, with some samples, including RS, showing a bimodal distribution. This variability could be due to the presence of sub-populations with transient high expression of these genes, resulting in an averaged increase in bulk RNA-Seq data compared to single-cell RNA-Seq. Supplemental Fig. 2D confirms that CI and SW cells were not senescent, as evidenced by their negative *β*-galactosidase staining.

### QASP Gene Expression Varies Across Cell Types and Quiescence Induction Methods

Using a similar approach as the one used to generate the QASP, we characterized genes upregulated in senescent cells to extract an unbiased list of SASP genes from the secretome. We combined the significant differentially expressed genes from our senescent scRNA-Seq data for LF1 cells across three different types of senescence (DDIS, OIS, RS) under low-oxygen conditions, with the bulk RNA-Seq data from senescent (RS) BJ and IMR90 cells generated under high oxygen used in our previous analysis. Additionally, we generated two time points of RS (Early senescence and Late senescence) in LF1 cells under low oxygen. From this pool of significantly expressed genes, we extracted those transcriptionally upregulated from the list of 1891 genes in the human secretome [13]. This subset of genes represents the transcriptional Senescence-Associated Secretory Phenotype (SASP), analogous to our previously generated QASP list but specific to senescence.

The greatest contribution to the QASP came from the high-oxygen dataset of BJ, IMR90, and Preadipocytes, followed by the low-oxygen IMR90 and single-cell LF1 quiescence datasets, with the lowest contribution observed in the low-oxygen LF1 dataset (Supplemental Fig. 3A left). On the other hand, the greatest contributor to the SASP gene list was the single-cell senescent dataset, followed by the low-oxygen LF1 dataset, with the lowest contribution being from the high-oxygen BJ and IMR90 senescent set (Supplemental Fig. 3A right). Gene Ontology (GO) analysis of all significantly upregulated QASP genes revealed enrichment of inflammatory pathways, including the Inflammatory Response, IL6-JAK-STAT3 signaling, and TNF*α* signaling via NF-*κ*B, among others (Supplemental Fig. 3B).

Surprisingly, the overlap between the QASP and the SASP was significant, with 205 genes unique to senescence, 159 genes unique to quiescence, and 589 genes shared between the two profiles (Fig. 3A and Supplemental Table 1). This overlap suggests that the inflammatory profile of quiescent cells may act as a precursor of the SASP, with quiescence representing an early stage that predisposes cells to transition into senescence.

**Figure 3.**
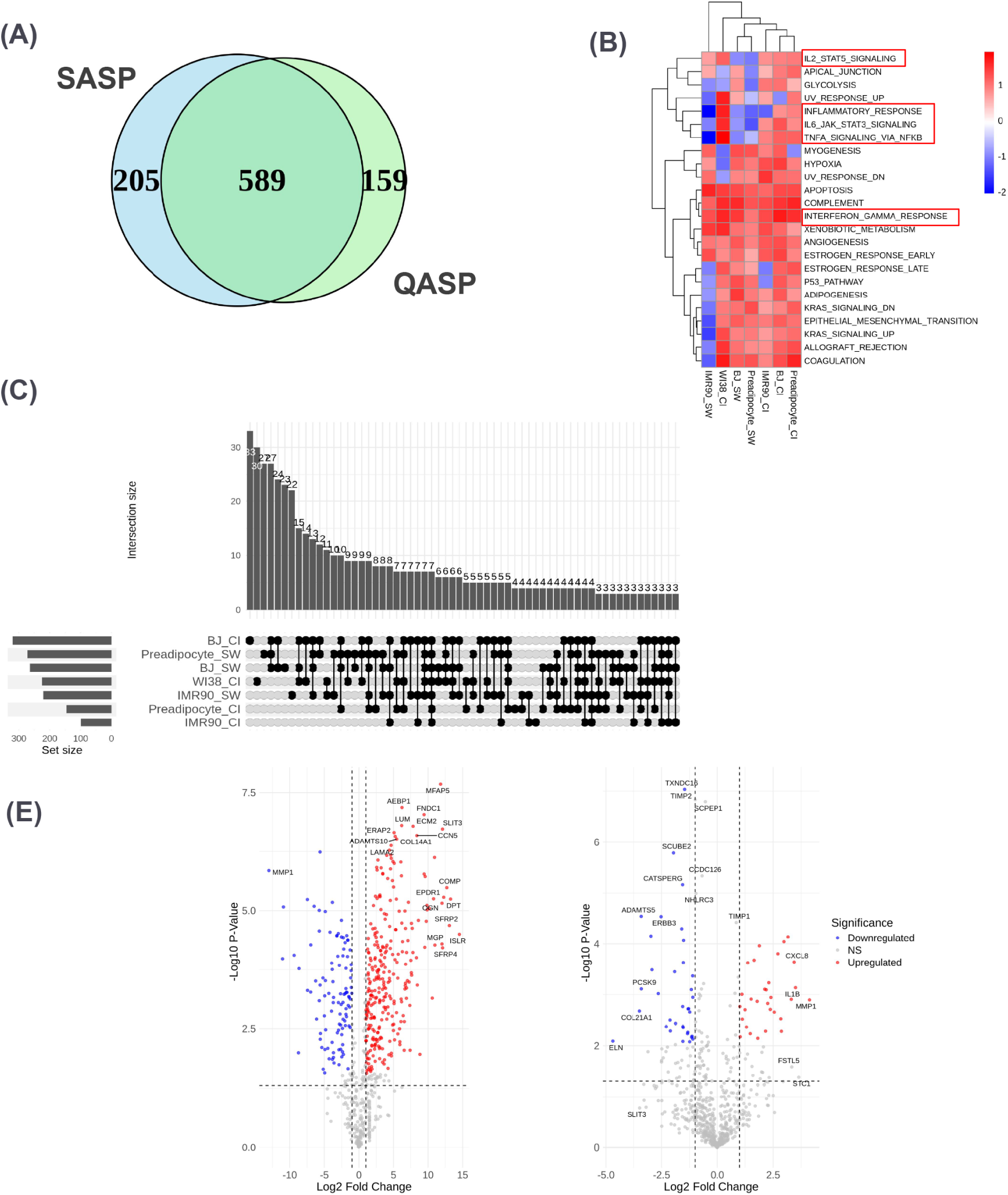
Comparison of QASP across cell lines and induction methods. (A) Venn diagram illustrating the overlap between SASP and QASP genes. (B) GSEA analysis of significantly upregulated pathways associated with QASP genes in quiescent versus proliferating cell comparisons (FDR *<* 0.05 and log_2_ FC *>* 1). Inflammatory pathways are highlighted. (C) UpSet plot showing overlap of significantly upregulated QASP genes (FDR *<* 0.05 and *|* log_2_ FC*| >* 1) across CI- and SW-induced quiescence in BJ, IMR90, WI38, and preadipocyte cell lines. Each vertical bar represents the number of QASP genes shared across the specified conditions. Only intersections containing three or more genes are displayed. (D) Volcano plots comparing quiescent versus proliferating cells (left) and CI versus SW (right), highlighting significantly differentially expressed QASP genes (FDR *<* 0.05 and log_2_ FC *>* 1) across cell lines. Upregulated genes are depicted in red and downregulated genes in blue.

To investigate the expression profile of QASP genes in high-oxygen conditions across quiescent (CI and SW) and senescent cells (RS) in BJ, IMR90 and preadipocytes, we filtered significant differentially expressed QASP genes across these experimental groups using the criteria of FDR *<* 0.05 and *|* log_2_ FC*| >* 1 (Supplemental Fig. 3C). In IMR90 cells, RS samples expressed QASP gene expression patterns similar pattern to those of CI and SW, although both quiescent comparisons were more similar to each other than to RS. A comparable pattern was observed in BJ cells, where the CI and SW comparisons clustered more closely together than RS, despite many genes showing upregulation or downregulation across all three conditions. In contrast, preadipocytes exhibited distinct differences in gene expression between CI vs. SW. Overall, each cell line demonstrated unique clustering, with quiescent CI and SW conditions grouping more closely than RS, and expression patterns differing between cell lines. This suggests that the QASP is both cell-line specific and influenced by the method of quiescence induction.

To assess the overlap of QASP genes across different cell lines and quiescence induction methods, we generated an UpSet plot based on the significantly upregulated QASP genes in each condition compared to proliferating controls (Fig. 3C). The overall overlap between all seven conditions was minimal, with only a small number of shared genes across combinations. BJ CI and Preadipocyte SW exhibited the largest overlap (33 genes), suggesting shared features in their transcriptional profiles. BJ CI also showed the highest number of unique QASP genes (30), followed by Preadipocyte SW (27). IMR90 cells had a low overlap between CI and SW. WI38 CI shared some overlap with other conditions, but did not dominate any intersection. This confirms that QASP signatures are highly context-specific, varying by both cell type and method of quiescence induction.

To further characterize the biological processes associated with the QASP, we performed GSEA analysis on the significant differentially expressed QASP genes in each cell line and quiescence induction method compared to proliferating (Fig. 3B). Interferon *gamma* and *alpha* response were upregulated in all comparisons, inflammatory response was upregulated in all comparisons except IMR90 SW, and TNF-*alpha* signaling was upregulated in BJ and preadipocyte cells but not IMR90. Gene Ontology (GO) term enrichment (Fig. 3B) of the QASP genes showed the most represented terms were those related to ECM organization, suggesting that quiescent cells may influence the physical properties of their microenvironment. Upregulation of chemotaxis, kinase signaling pathways and leukocyte migration suggests that quiescent cells might play a role in signaling to immune cells or surrounding cells via inflammatory processes.

To understand the differential expression of QASP genes in quiescent cells, we generated volcano plots comparing quiescent cells (combined CI and SW) to proliferating cells (Fig. 3E, left), as well as CI-induced versus SW-induced quiescent cells (Fig. 3E, right). Among the genes significantly upregulated in quiescent cells relative to proliferating cells, we observed enrichment of extracellular matrix (ECM)-associated and matricellular proteins, including *AEBP1*, *LUM*, *COL14A1*, *MFAP5*, *ADAMTS10*, and *CCN5*, highlighting a coordinated ECM remodeling response during quiescence (Table 2). In contrast, the matrix-degrading enzyme *MMP1* was significantly downregulated, suggesting suppression of ECM turnover in the quiescent state. When comparing CI- and SW-induced quiescence, we observed that pro-inflammatory cytokines such as *IL1B* and *CXCL8* were upregulated in CI-induced cells, while ECM-related genes including *ELN*, *COL21A1*, and *ADAMTS5* were more highly expressed in SW-induced cells (Table 3). This suggests that the method of quiescence induction differentially modulates inflammatory and structural remodeling signatures.

**Table 2:** Top QASP genes significantly up-or downregulated in Quiescent vs Proliferative cells.

| **Gene** | **Condition** | **Function** |
|---|---|---|
| AEBP1 | Upregulated | Transcriptional repressor involved in adipogenesis and smooth muscle differentiation. |
| LUM | Upregulated | Proteoglycan involved in collagen fibrillogenesis and tissue repair. |
| COL14A1 | Upregulated | Fibril-associated collagen that regulates fibrillogenesis. |
| ERAP2 | Upregulated | Aminopeptidase involved in antigen processing for MHC class I. |
| MFAP5 | Upregulated | Glycoprotein contributing to elastic fiber formation. |
| LAMA2 | Upregulated | Laminin subunit contributing to basement membrane stability. |
| ADAMTS10 | Upregulated | Metalloprotease involved in ECM organization. |
| FNDC1 | Upregulated | Fibronectin-like domain protein involved in adhesion. |
| ECM2 | Upregulated | ECM protein promoting cell-matrix interactions. |
| CCN5 | Upregulated | Matricellular signaling modulator for proliferation and migration. |
| SLIT3 | Upregulated | Guidance cue involved in cell migration and tissue organization. |
| MMP1 | Downregulated | Collagenase involved in ECM breakdown. |
| COMP | Upregulated | Cartilage ECM structural protein. |
| EPDR1 | Upregulated | Protein involved in cell adhesion and matrix remodeling. |
| DPT | Upregulated | Dermatopontin; supports ECM structure and signaling. |
| OGN | Upregulated | Regulator of collagen fibril organization and bone growth. |
| SFRP2 | Upregulated | Wnt signaling pathway modulator. |
| ISLR | Upregulated | Immunoglobulin-like protein possibly involved in regeneration. |
| MGP | Upregulated | Inhibitor of vascular calcification. |
| SFRP4 | Upregulated | Wnt signaling pathway modulator. |

**Table 3:** Top QASP genes significantly changed between Contact-Inhibited (CI) and Serum-Withdrawn (SW) quiescent states.

| **Gene** | **Condition** | **Function** |
|---|---|---|
| TXNDC16 | Downregulated | Thioredoxin-domain protein involved in redox homeostasis. |
| TIMP2 | NS | Inhibitor of matrix metalloproteinases; regulates ECM turnover. |
| SCPEP1 | NS | Serine carboxypeptidase involved in protein catabolism. |
| SCUBE2 | Downregulated | Cell adhesion molecule with signaling roles. |
| CCDC126 | NS | Poorly characterized; coiled-coil domain-containing protein. |
| NHLRC3 | NS | RING-finger protein; function not fully characterized. |
| CATSPERG | Downregulated | Possible ion channel regulator; function unknown. |
| ADAMTS5 | Downregulated | Metalloprotease involved in cartilage matrix degradation. |
| ERBB3 | Downregulated | Receptor tyrosine kinase involved in cell signaling. |
| TIMP1 | NS | Inhibits ECM-degrading metalloproteinases. |
| CXCL8 | Upregulated | Chemokine IL-8; promotes neutrophil recruitment. |
| PCSK9 | Downregulated | Regulates LDL receptor turnover and cholesterol homeostasis. |
| IL1B | Upregulated | Pro-inflammatory cytokine, key immune mediator. |
| MMP1 | Upregulated | ECM-degrading enzyme (collagenase). |
| COL21A1 | Downregulated | Structural collagen protein in ECM. |
| ELN | Downregulated | Encodes elastin; key structural ECM component. |
| FSTL5 | NS | Secreted glycoprotein; function remains unclear. |
| STC1 | NS | Hormone involved in calcium/phosphate regulation. |
| SLIT3 | NS | Guidance molecule in tissue patterning and remodeling. |

### Oxygen Levels Modulate QASP Gene Expression in Quiescent Cells

Mammalian cells are commonly cultured under hyperoxic conditions. While most cells experience oxygen levels ranging from 1% to 6% *in vivo* (physioxia), nearly all mammalian cell culture is conducted in humidified atmospheric air (18-20%) at 37°C with 5% CO_2_ added [24]. Oxygen participates in many metabolic reactions, some of which are sensitive to Oxygen levels between physiological and the approximately 18% found in standard incubators. Elevated oxygen in cell cultures can increase reactive oxygen species (ROS), contributing to changes in cellular senescence, differentiation and apoptosis [25].

To explore the effects of oxygen in quiescence induction, we cultured IMR90 cells under either 2.5% O_2_ (low oxygen) or atmospheric O_2_ (high oxygen) and induced quiescence through CI or SW. We also induced DDIS with etoposide treatment for two weeks under low and high oxygen. Quantitative PCR (qPCR) was performed on cell cycle inhibitor and senescence marker *CDKN2A* and inflammatory genes *IL1B, IL6* and *IL8* to assess whether their expression was affected by oxygen levels (Supplemental Fig. 4A). As expected, *CDKN2A* expression levels were higher in senescent cells, with SW showing slightly higher levels than CI and overall expression patterns remained consistent across oxygen conditions. *IL1B* exhibited the greatest difference in expression between low and high oxygen, with *IL1B* levels being higher in CI compared to senescent cells under low oxygen. Additionally, *IL6* showed increased levels in SW under low oxygen, while *IL8* was elevated in CI under low oxygen compared to senescence. These results demonstrate that oxygen levels influence the expression of inflammatory genes during quiescence and senescence.

To further explore these effects, we performed bulk RNA-Seq on the quiescent samples induced by either CI or SW under both low and high-oxygen conditions (Fig. 4A). PCA revealed that proliferating cells clustered along the PC1 axis while the quiescent cells displayed less distinction based on induction method and oxygen levels. Specifically, CI samples tended to cluster slightly more to the left, whereas SW samples clustered more to the right along PC1. Greater variability was observed along PC2, with SW samples under low and high-oxygen conditions clustering closer together, while CI samples under high and low-oxygen conditions clustered on opposite sides of the graph, indicating increased gene expression variability under high-oxygen conditions (Supplemental Fig. 4B).

**Figure 4.**
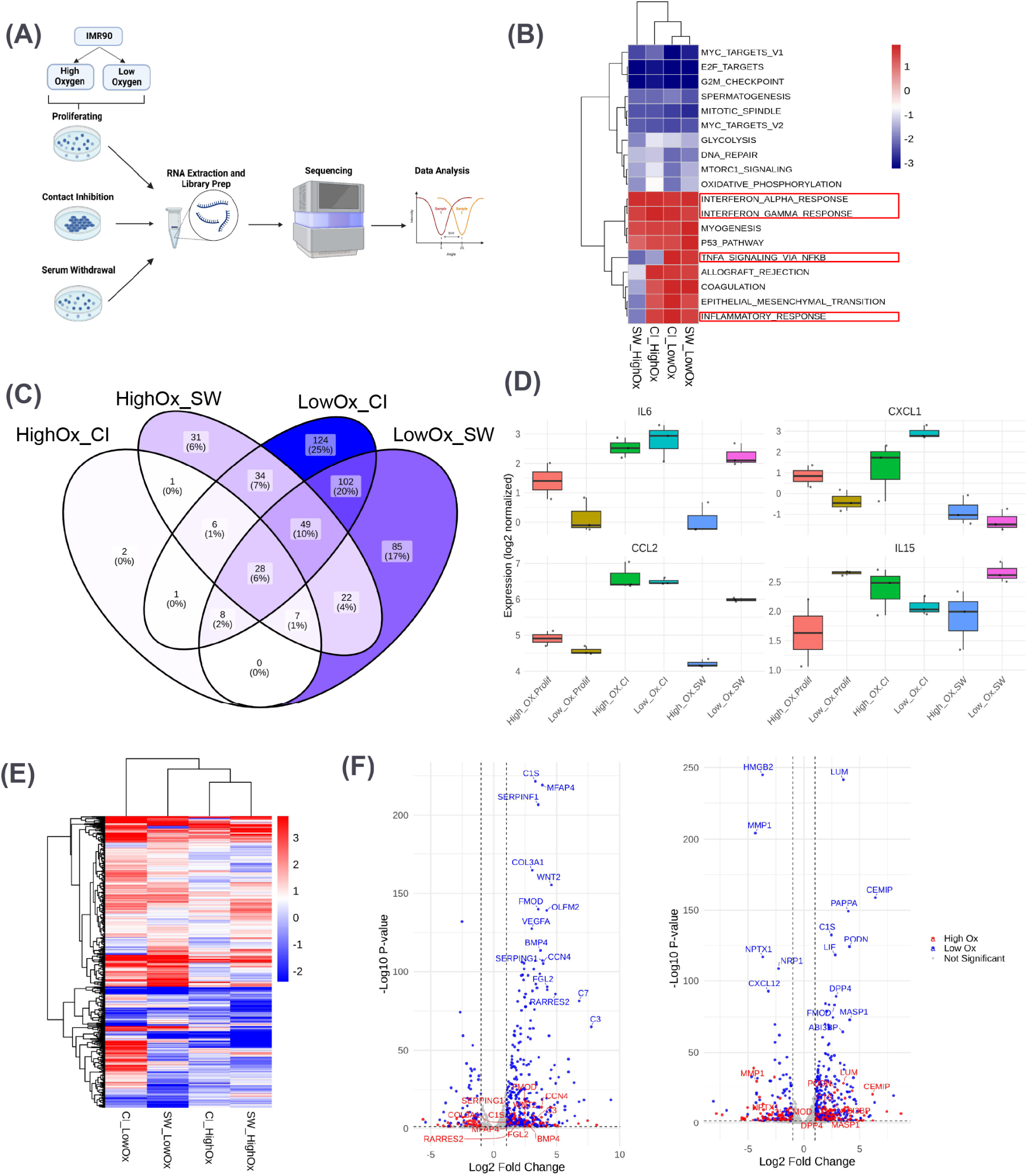
Comparative analysis of quiescence induction under high- and low-oxygen conditions in IMR90 cells. (A) Schematic representation of the experimental design, illustrating IMR90 cells cultured under high-oxygen (20%) and low-oxygen (2.5%) conditions for each state: proliferation, contact inhibition, and serum withdrawal. (B) GSEA heatmap depicting significantly differentially upregulated and downregulated pathways (FDR *<* 0.05 and log_2_ FC *>* 1) in IMR90 cells across oxygen levels and induction methods compared with proliferating cells. Inflammatory pathways are highlighted. (C) Venn diagram illustrating the overlap of differentially expressed QASP genes (FDR *<* 0.05 and log_2_ FC *>* 1) between oxygen conditions and induction methods compared with proliferating cells. (D) Box plots showing expression of inflammatory genes *IL6*, *CXCL1*, *CCL2*, and *IL15* in CI, SW, and proliferating IMR90 or LF1 cells cultured under high-versus low-oxygen conditions. (E) Heatmap showing significant upregulation and downregulation of QASP genes (FDR *<* 0.05 and log_2_ FC *>* 1) across induction methods under both oxygen levels compared with proliferating cells. (F) Volcano plots depicting differential gene expression under high-oxygen (red) and low-oxygen (blue) conditions for CI versus Prolif (left) and SW versus Prolif (right). Highlighted genes include the top 10 genes by log_2_ fold change and the top 10 by *−* log_10_ p-value among QASP genes.

A heatmap of differential gene expression analysis showed clustering by oxygen levels and not by induction method, with the different oxygen levels showing similar expression patterns, while there was a difference in expression between CI and SW, particularly at low oxygen. This supports our findings that oxygen influences quiescence in a manner dependent on the induction method (Supplemental Fig. 4C). GSEA analysis showed consistent upregulation of interferon *alpha* and *gamma* responses across all conditions, while TNF*α* signaling via NF*κ*B was only upregulated in low oxygen and the inflammatory response was upregulated in all conditions except SW under high oxygen (Fig. 4B). This analysis revealed that oxygen levels significantly influence the expression of QASP genes and associated pathways during quiescence induction.

Next, we wanted to explore whether different quiescence induction methods affected the expression of cell cycle inhibitors under high vs. low-oxygen conditions (Supplemental Fig. 4D). Violin plots of our bulk RNA-Seq data revealed that *CDKN1A, CDKN2A* and *CDKN2B* expression was decreased in low oxygen, although expression under low oxygen showed higher variability, reflecting a heterogeneous cellular response to reduced oxygen. Expression of *CDKN1B* and *CKDN2C* was higher under low oxygen, while expression of *CDKN2D* and *CDKN1C* were slightly higher but did not show marked differences under low vs. high oxygen. These findings suggest that the expression of certain cell cycle inhibitors may be stabilized or upregulated in response to hypoxia.

To investigate the influence of oxygen on the expression of QASP genes, we performed differential gene expression analysis of QASP genes under high vs. low oxygen. We observed pronounced differences in secretory expression between high and low-oxygen conditions, with CI under low oxygen showing the most substantial difference between the two oxygen level conditions. It should be noted that the comparisons clustered by oxygen levels and not induction method, which is indicative of the effect of oxygen in QASP gene expression (Fig. 4E). A Venn diagram (Fig. 4C)showed that only 28 QASP gene overlapped across the four conditions (Table 4. Low oxygen CI and SW accounted for the majority of QASP genes, with the largest overlap, consisting of 102 genes (21%). In contrast, CI and SW under high oxygen only shared one gene. Additionally, SW samples under both low and high-oxygen conditions had an overlap of 49 genes, while CI samples across high and low oxygen showed a small overlap of 5 genes. Expression of inflammatory genes *IL6, CXCL1, CCL2* and *IL15* showed upregulation more consistent across induction method than oxygen levels, with CI showing the most consistent upregulation (Fig. 4D).

**Table 4:**
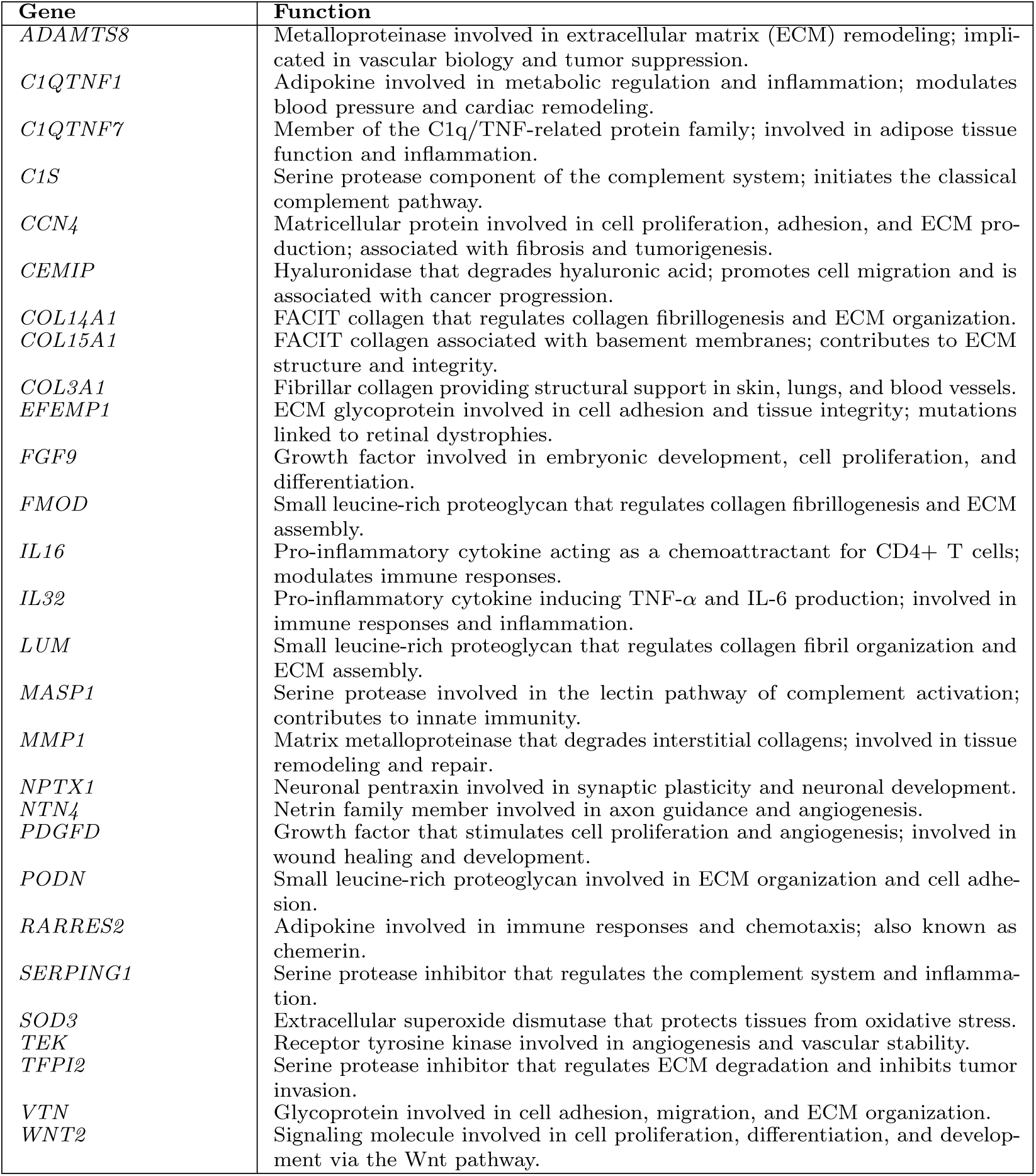
Functional Annotations of Overlapping QASP Genes.

| **Gene** | **Function** |
|---|---|
| *ADAMTS8* | Metalloproteinase involved in extracellular matrix (ECM) remodeling; impli-cated in vascular biology and tumor suppression. |
| *C1QTNF1* | Adipokine involved in metabolic regulation and inflammation; modulates blood pressure and cardiac remodeling. |
| *C1QTNF7* | Member of the C1q/TNF-related protein family; involved in adipose tissue function and inflammation. |
| *C1S* | Serine protease component of the complement system; initiates the classical complement pathway. |
| *CCN4* | Matricellular protein involved in cell proliferation, adhesion, and ECM pro-duction; associated with fibrosis and tumorigenesis. |
| *CEMIP* | Hyaluronidase that degrades hyaluronic acid; promotes cell migration and is associated with cancer progression. |
| *COL14A1* | FACIT collagen that regulates collagen fibrillogenesis and ECM organization. |
| *COL15A1* | FACIT collagen associated with basement membranes; contributes to ECM structure and integrity. |
| *COL3A1* | Fibrillar collagen providing structural support in skin, lungs, and blood vessels. |
| *EFEMP1* | ECM glycoprotein involved in cell adhesion and tissue integrity; mutations linked to retinal dystrophies. |
| *FGF9* | Growth factor involved in embryonic development, cell proliferation, and differentiation. |
| *FMOD* | Small leucine-rich proteoglycan that regulates collagen fibrillogenesis and ECM assembly. |
| *IL16* | Pro-inflammatory cytokine acting as a chemoattractant for CD4+ T cells; modulates immune responses. |
| *IL32* | Pro-inflammatory cytokine inducing TNF-α and IL-6 production; involved in immune responses and inflammation. |
| *LUM* | Small leucine-rich proteoglycan that regulates collagen fibril organization and ECM assembly. |
| *MASP1* | Serine protease involved in the lectin pathway of complement activation; contributes to innate immunity. |
| *MMP1* | Matrix metalloproteinase that degrades interstitial collagens; involved in tissue remodeling and repair. |
| *NPTX1* | Neuronal pentraxin involved in synaptic plasticity and neuronal development. |
| *NTN4* | Netrin family member involved in axon guidance and angiogenesis. |
| *PDGFD* | Growth factor that stimulates cell proliferation and angiogenesis; involved in wound healing and development. |
| *PODN* | Small leucine-rich proteoglycan involved in ECM organization and cell adhe-sion. |
| *RARRES2* | Adipokine involved in immune responses and chemotaxis; also known as chemerin. |
| *SERPING1* | Serine protease inhibitor that regulates the complement system and inflamma-tion. |
| *SOD3* | Extracellular superoxide dismutase that protects tissues from oxidative stress. |
| *TEK* | Receptor tyrosine kinase involved in angiogenesis and vascular stability. |
| *TFPI2* | Serine protease inhibitor that regulates ECM degradation and inhibits tumor invasion. |
| *VTN* | Glycoprotein involved in cell adhesion, migration, and ECM organization. |
| *WNT2* | Signaling molecule involved in cell proliferation, differentiation, and develop-ment via the Wnt pathway. |

We generated volcano plots to compare QASP genes differentially expressed in CI vs. proliferating (Fig. 4F left) and SW vs. proliferating cells (Fig. 4F right) highlighting the differences between high and low-oxygen conditions. We observed significant differences in QASP genes upregulated and downregulated in each induction method, with oxygen level significantly impacting gene expression. Low-oxygen conditions displayed the most pronounced differences, reflected in both the highest fold change and p-value rankings.

Given the substantial role that oxygen plays in cellular metabolism and gene expression, we specifically focused on IMR90 and LF1 cells under low-oxygen conditions to more closely approximate the physiological environments these cells would encounter *in vivo*. By limiting oxygen levels to 2.5%, we aimed to minimize the potential artifacts associated with high-oxygen culture conditions. In IMR90 cells, quiescence was induced via CI and SW, while in LF1 cells, quiescence was induced via SW. Additionally, we analyzed two RS time points in LF1 cells: early senescence and late senescence. Batch-corrected PCA showed proliferating cells clustered toward the right along PC1, CI and SW clustered in the middle, and early and late senescence on the left (Fig. 5A). Non-batch corrected data can be observed in Supplemental Fig.5A. A heatmap of significant differentially expressed genes showed clustering and similar expression patters by cell line, with SW for LF1 showing a slight difference in gene expression patterns compared to LF1 senescence (both early and late sen), which were very similar to each other (Supplemental Fig. 5B).

**Figure 5.**
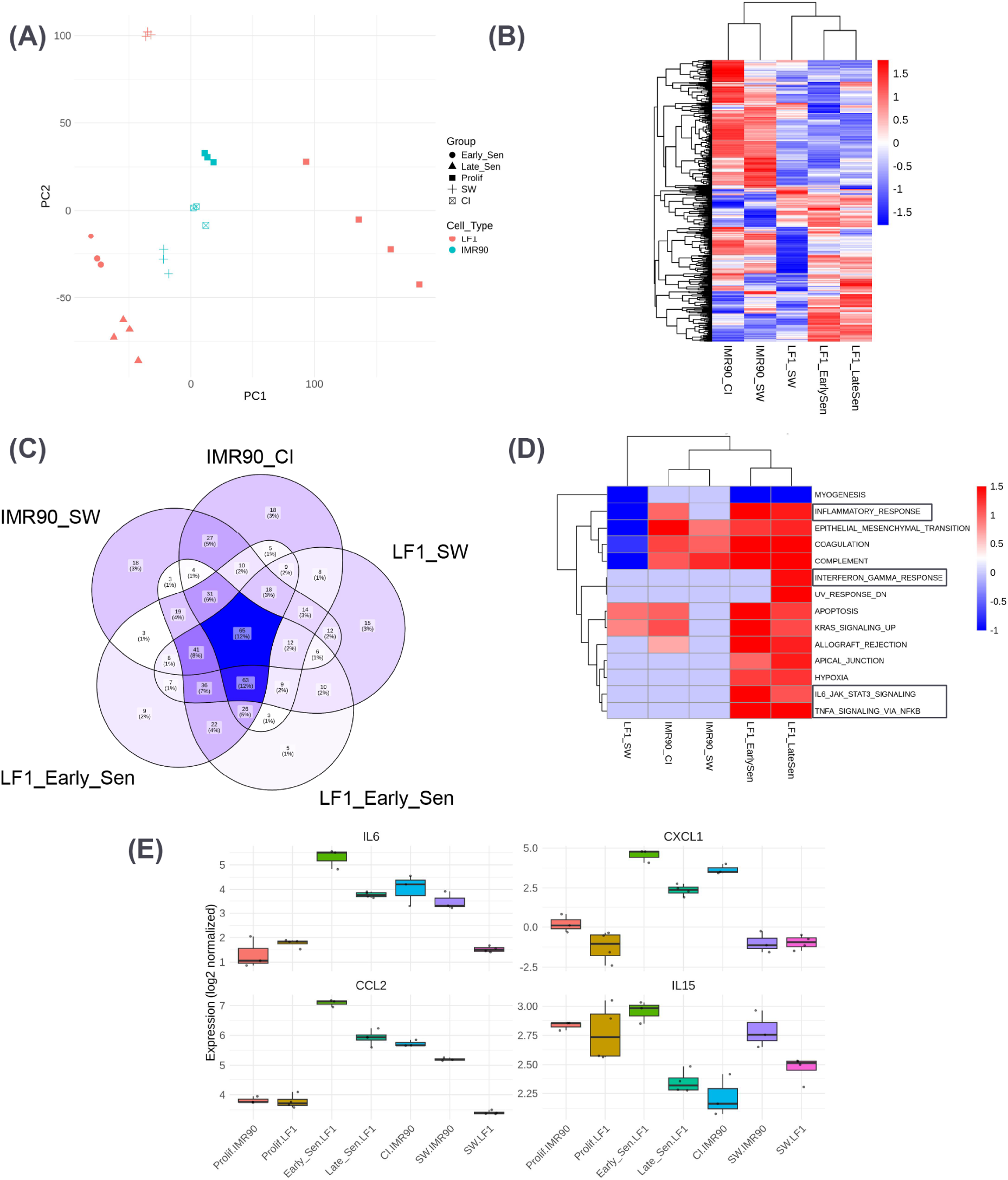
Comparative analysis of quiescent and senescent LF1 cells and IMR90 cells under low-oxygen conditions. (A) PCA plot demonstrating the separation of samples by both cell type and condition after batch correction. LF1 cells are depicted in blue and IMR90 cells in red. (B) Heatmap illustrating significantly differentially expressed QASP genes (FDR *<* 0.05 and log_2_ FC *>* 1) across all contrasts under low-oxygen conditions: IMR90 CI versus Prolif, IMR90 SW versus Prolif, LF1 SW versus Prolif, Early Sen versus Prolif, and Late Sen versus Prolif. (C) Venn diagram showing the overlap of significantly differentially expressed QASP genes (FDR *<* 0.05 and log_2_ FC *>* 1) across comparisons between conditions and proliferating cells. (D) GSEA heatmap displaying differentially upregulated and downregulated pathways for significantly differentially expressed QASP genes (FDR *<* 0.05 and log_2_ FC *>* 1) across conditions under low oxygen compared with proliferating cells. Inflammatory pathways are highlighted. (E) Box plots showing expression of inflammatory genes *IL6*, *CXCL1*, *CCL2*, and *IL15* in quiescent or senescent LF1 cells cultured under low-oxygen (2.5%) conditions.

To investigate the expression profile of QASP genes in low oxygen, quiescent (CI and SW) and senescent cells (RS) for IMR90 and LF1 cells, we filtered significantly expressed QASP genes across these comparisons using criteria of FDR *<* 0.05 and log_2_FC *>* 1. Quiescent IMR90 cells exhibited a more consistent pattern of QASP upregulation and downregulation, while LF1 SW showed the least amount of QASP upregulation, in a pattern distinct from early and late senescence. While comparisons clustered by cell line, both SW from IMR90 and LF1 clustered closer together, suggesting a similar pattern of expression for this induction method. As expected, senescent cells clustered together as well (Fig. 5B). A Venn diagram (Fig. 5C) revealed a 13% overlap of QASP genes across all conditions, with IMR90 CI vs. proliferating overlapping by 12% with both LF1 senescent conditions. This overlap reflected a more inflammatory phenotype in CI-induced cells compared to SW-induced cells under low oxygen.

GSEA of significantly differentially expressed genes indicated that SW conditions for both IMR90 and LF1 were more similar to each other and clustered together, while CI clustered closer to senescent cells. Notably, IMR90 CI vs. proliferating and both senescent LF1 comparisons showed upregulation of TNF-*α* signaling via NF-*κ*B, interferon *gamma* and inflammatory response pathways. Only senescent cells showed upregulation of IL6 JAK STAT3 signaling pathway (Fig. 5D). Violin plots depicting the expression levels of inflammatory genes *IL6, CXCL1, CCL2* and *IL15* showed the most upregulation in senescence, followed by CI IMR90 and to a lesser extent, in SW IMR90. LF1 SW did not exhibit upregulation of these genes (Supplemental Fig. 6C. The reduced expression observed under low-oxygen conditions suggests that culturing cells at physiological oxygen levels may suppress the QASP and related inflammatory responses in a cell line dependent manner.

To better understand the relationship between QASP expression in quiescence and senescence, we isolated upregulated QASP genes shared across SW vs. proliferating and both senescent comparisons in LF1 cells. The final list consisted of 34 genes, which we categorized based on their functions. Most of these genes were more highly expressed in early senescence than in quiescence and late senescence. Specifically, quiescent cells showed more upregulation of genes involving cell adhesion and migration, while both senescence comparisons demonstrated increased expression of genes related to ECM and tissue remodeling, as well as growth factors and receptor signaling immune modulation and inflammation (Fig. 5E). Overall, under low-oxygen conditions, SW-induced quiescence did not appear to display an inflammatory profile, while CI-induced quiescence and senescence did.

Our findings demonstrate that oxygen levels significantly influence the QASP expression of quiescent cells, with distinct patterns emerging under high versus low oxygen. Low oxygen, which better approximates physiological conditions, suppressed inflammatory responses associated with QASP expression, particularly in SW-induced quiescence. In contrast, CI-induced quiescence retained a more inflammatory phenotype. This translated to the pathways upregulated under these conditions as well, with quiescent cells showing upregulation of interferon pathways, with SW showing a less inflammatory profile than CI. These findings suggest that low oxygen may attenuate inflammatory responses, offering insights into how oxygen availability may influence quiescent cells and their transition into senescence.

### Quiescent Cells Actively Secrete Inflammatory Cytokines

While transcriptional analysis revealed an inflammatory signature in quiescent cells, it remained unclear whether this profile translated into increased secretion of cytokines. To assess the functional relevance of the QASP and confirm whether quiescent cells actively release pro-inflammatory factors, we performed a multiplexed cytokine assay on conditioned media from IMR90 fibroblasts induced into quiescence via CI and SW and compared them to proliferating and senescent (etoposide-treated) controls cultured in high oxygen. We also induced quiescence via CI and SW and compared them to proliferating under low oxygen conditions.

Analysis of cytokine secretion revealed an induction method-dependent inflammatory profile in quiescent cells. Several cytokines, including *IL6*, *IL1B*, *IL15*, and *CCL2*, were secreted at significantly higher levels in CI-induced quiescent cells compared to both proliferating and SW-induced quiescent cells. This mirrors the induction method-specific transcriptional regulation observed in the QASP, where CI-induced quiescent cells exhibited a stronger pro-inflammatory signature than SW-induced quiescent cells. Notably, IL6 and CCL2 secretion in CI quiescent cells exceeded the assay’s maximum limit of detection, whereas proliferating cells had readings at the lower limit of detection, precluding statistical significance despite substantial fold increases in both high oxygen (Figure 6 (A)) and low oxygen (Figure 6 (B))

**Figure 6.**
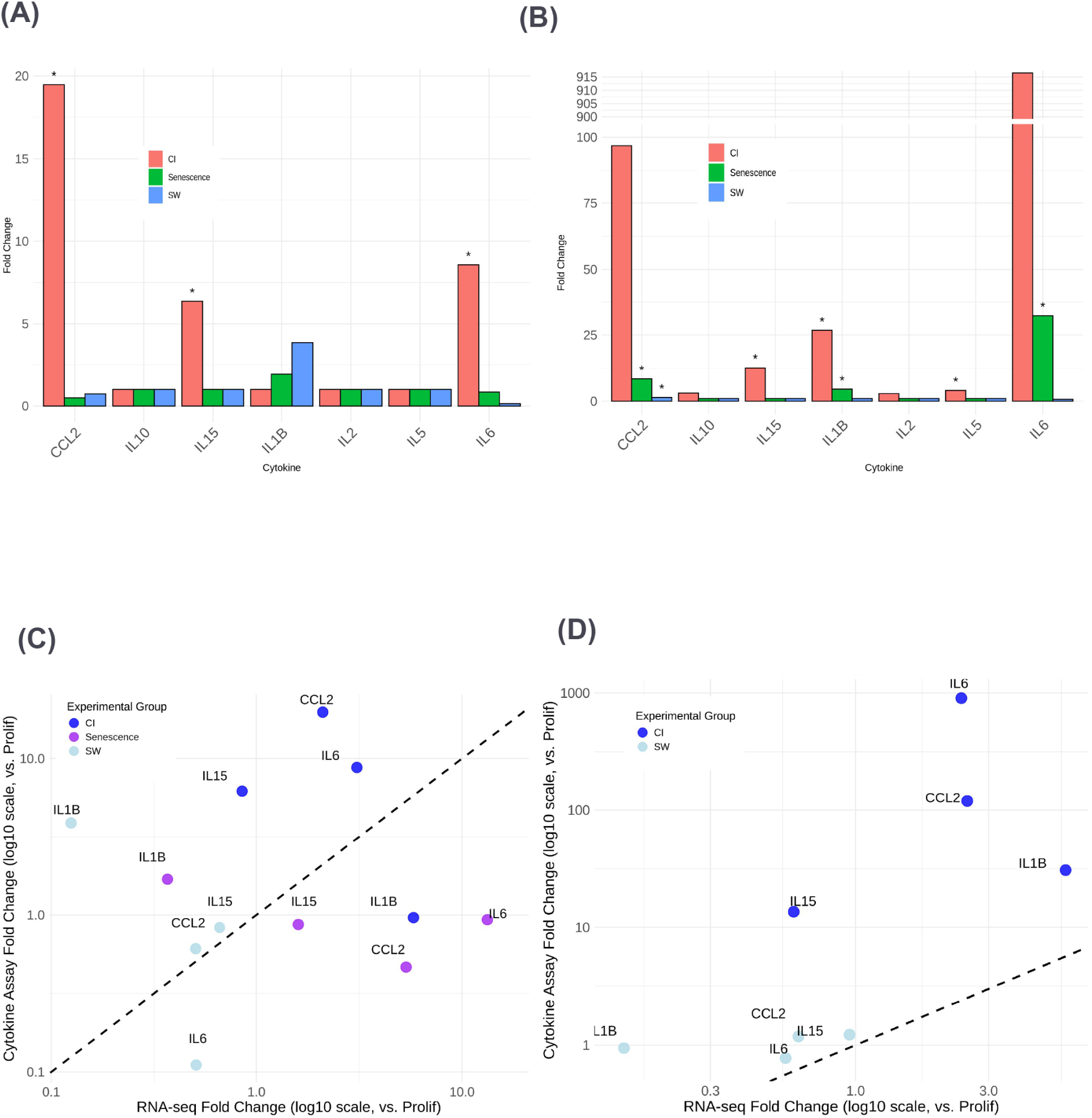
Analysis of cytokine secretion in quiescent and senescent IMR90 cells compared with proliferating cells under high- and low-oxygen conditions. (A) Box plots displaying cytokine secretion fold changes relative to proliferating cells for quiescence induced via contact inhibition (CI), serum withdrawal (SW), and etoposide-induced senescence for IL-10, IL-15, IL-1*β*, IL-2, IL-5, IL-6, and CCL2 under high-oxygen conditions. Experimental groups are shown as follows: CI in red, SW in blue, and senescence in green. Statistically significant differences (*p <* 0.05, one-sided one-sample *t*-test) are indicated by asterisks. For IL-6 and CCL2, statistical significance could not be formally determined because of data truncation at the assay detection limits. In CI-induced quiescence, cytokine levels exceeded the upper limit of quantification (ULOQ), whereas in proliferating cells, values approached the lower limit of quantification (LLOQ). This loss of dynamic range prevented accurate p-value calculation, although the large fold changes suggest substantial differences in cytokine secretion. (B) Box plots displaying cytokine secretion fold changes relative to proliferating cells for quiescence induced via contact inhibition (CI) and serum withdrawal (SW) for IL-10, IL-15, IL-1*β*, IL-2, IL-5, IL-6, and CCL2 under low-oxygen conditions. Experimental groups are shown as follows: CI in red and SW in blue. Statistically significant differences (*p <* 0.05, one-sided one-sample *t*-test) are indicated by asterisks. For IL-6 and CCL2, statistical significance could not be formally determined because of data truncation at the assay detection limits. In CI-induced quiescence, cytokine levels exceeded the upper limit of quantification (ULOQ), whereas in proliferating cells, values approached the lower limit of quantification (LLOQ). This loss of dynamic range prevented accurate p-value calculation, although the large fold changes suggest substantial differences in cytokine secretion. (C) Scatter plot comparing RNA-seq fold changes (x-axis) and cytokine secretion fold changes (y-axis) under high-oxygen conditions. A dashed diagonal line represents the 1:1 relationship, indicating proportional changes in cytokine secretion and gene expression. Contact inhibition (CI) is denoted in blue, serum withdrawal (SW) in light blue, and senescence in purple. (D) Scatter plot comparing RNA-seq fold changes (x-axis) and cytokine secretion fold changes (y-axis) under low-oxygen conditions. A dashed diagonal line represents the 1:1 relationship. Contact inhibition (CI) is denoted in blue and serum withdrawal (SW) in light blue.

To determine whether transcriptional upregulation of QASP genes correlates with active cytokine secretion, we compared RNA-seq fold changes with cytokine levels. Scatter plot analysis revealed a strong correlation between transcriptional and secreted cytokine levels, with most quiescence-associated inflammatory genes exhibiting increased secretion relative to proliferating cells in high oxygen (Figure 6 (C) and low oxygen (Figure 6 (D). However, the extent of cytokine secretion was not always directly proportional to gene expression, suggesting additional layers of post-transcriptional regulation. Specifically, CI-induced quiescent cells exhibited disproportionately higher cytokine levels relative to RNA expression for *IL6*, *IL1B*, *IL15*, and *CCL2*, indicating potential post-translational modifications that enhance secretion. Conversely, senescent cells in high oxygen displayed higher transcriptional levels with comparatively lower cytokine secretion, suggesting possible post-translational repression mechanisms.

These findings establish the QASP as a functional inflammatory program, extending beyond transcriptional changes to active cytokine secretion. Our results support the notion that quiescent cells actively contribute to the inflammatory microenvironment, potentially priming surrounding cells for senescence and tissue remodeling.

### QASP Expression Is Elevated in Fibroblasts and Myofibroblasts in Idiopathic Pulmonary Fibrosis

Fibroblasts are vital for maintaining tissue integrity and play a key role in tissue support by producing and remodeling the ECM. Upon injury, fibroblasts can become activated and differentiate into myofibroblasts; cells characterized by enhanced contractile capacity and increased ECM production [26]. Persistent fibroblast activation, driven by the prolonged exposure to pro-inflammatory factors, can create a feedback loop that disrupts normal repair and leads to chronic wounds, fibrosis, and scarring [27]. Idiopathic Pulmonary Fibrosis (IPF) is a chronic lung disease characterized by pathological epithelial remodeling and accumulation of ECM [28]. In IPF, myofibroblasts are thought to arise through cytokine-induced differentiation of fibroblasts [29].

To investigate whether the QASP expression observed in our *in vitro* fibroblast studies is also evident in healthy vs. fibrotic lung tissue, we analyzed a publicly available scRNA-Seq dataset of human lung tissue [28]. Our analysis focused on fibroblast populations classified as either “healthy” or “IPF” (Fig. 7A). While the proportions proliferating and quiescent cells were similar between healthy and IPF fibroblasts, a higher proportion of senescent cells was observed in IPF (Fig. 7B). Cells formed distinct clusters based on proliferating, quiescent or senescent states (Supplemental Fig. 7A). For subsequent analysis, we restricted our focus to quiescent fibroblasts. These cells did not cluster by health condition, though each cluster contained subsets of healthy or IPF cells (Supplemental Fig. 7B).

**Figure 7.**
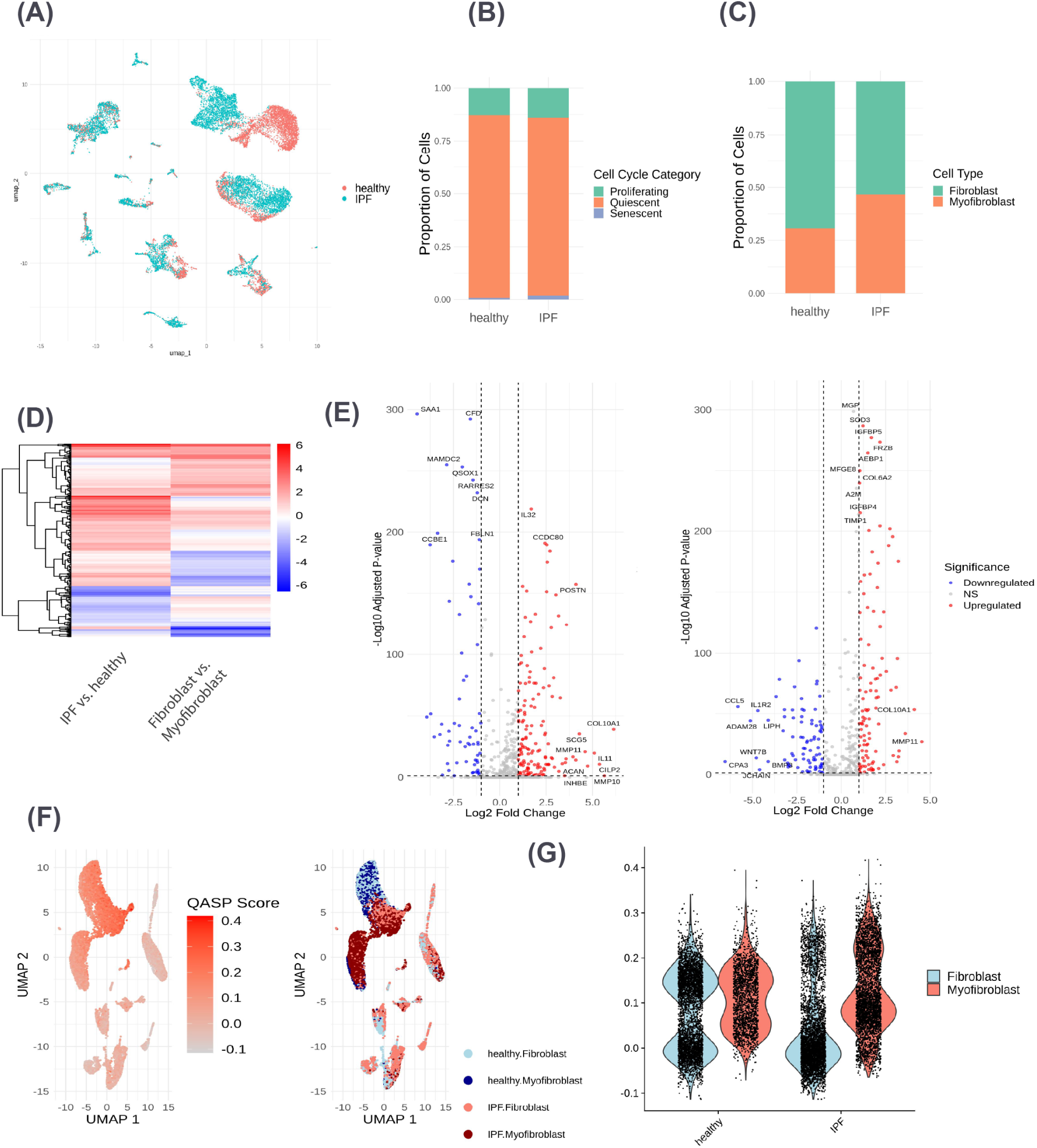
Comparative analysis of QASP in fibroblasts from healthy and IPF lung tissue using scRNA-seq data. (A) Uniform Manifold Approximation and Projection (UMAP) plots showing healthy versus IPF cells across all fibroblast populations (proliferating, quiescent, and senescent), illustrating that cells do not cluster by health condition. (B) Bar plot quantifying the proportions of proliferating, quiescent, and senescent fibroblasts in healthy versus IPF samples. Proliferating cells are shown in green, quiescent cells in orange, and senescent cells in blue. (C) Bar plot comparing the proportions of fibroblasts and myofibroblasts in healthy versus IPF samples. (D) Heatmap illustrating significantly differentially expressed QASP genes (FDR *<* 0.05 and log_2_ FC *>* 1) across IPF versus healthy and fibroblasts versus myofibroblasts. (E) Volcano plots demonstrating the upregulation and downregulation of QASP genes in myofibroblasts versus fibroblasts (left) and IPF versus healthy cells (right). Highlighted genes include the top 10 by log_2_ fold change and the top 10 by *−* log_10_ adjusted p-value. Upregulated genes are depicted in red and downregulated genes in blue. (F) UMAP plots showing QASP scores in quiescent fibroblasts (left) compared with their health condition (healthy versus IPF) and cell type (fibroblast versus myofibroblast) (right). Cells with higher QASP module scores cluster together toward the top of the UMAP. (G) Violin plots showing QASP module scores comparing fibroblasts and myofibroblasts in healthy and IPF samples.

To assess fibroblasts and myofibroblasts proportions in healthy vs. IPF tissue, we classified myofibroblasts using a set of markers (See Methods Section 5.8 and Supplemental Fig. 7C). Violin plots of additional myofibroblast markers (not included in the initial classification) revealed significantly higher expression levels in IPF compared to healthy tissue (Supplemental Fig. 7D). While the fibroblast proportions were similar between healthy vs. IPF tissue, the proportion of myofibroblasts was higher in IPF (Fig. 6C). This trend held across proliferating, quiescent and senescent cells in IPF, with a higher proportion of myofibroblasts observed in each state (Supplemental Fig. 7E).

We filtered significantly differentially expressed QASP genes in IPF vs. healthy quiescent fibroblasts and found distinct sets of upregulated genes in each condition (Fig. 7D). Comparing QASP expression between myofibroblasts and fibroblasts revealed an even more pronounced difference, with many QASP genes showing notably higher expression levels in myofibroblasts than in fibroblasts (Fig. 7E left).When we examined the significantly expressed QASP genes across IPF vs. healthy and myfibroblasts vs. fibroblasts comparisons, we observed that many upregulated genes were shared. However, some QASP genes were upregulated in one comparison and downregulated in the other (Fig. 7E right).

To further investigate the differences in QASP expression between fibroblasts and myofibroblasts in healthy vs. IPF tissue, we created a QASP module score and applied it to the differentially expressed genes in both comparisons. (Fig. 7F). We found that cells cluster not by cell type or disease state but by inflammatory levels (determined by the QASP score). Using the module score, we classified the cells as either “High QASP” or “Low QASP” and conducted differential gene expression between these groups. We observed a bimodal distribution of QASP expression, especially in healthy tissue, with IPG myofibroblasts showing the highest QASP module score (Fig. 7G). GO term analysis of the significantly differentially expressed QASP genes highlighted distinct pathway enrichments between fibroblasts and myofibroblasts. Fibroblasts showed upregulation in immune response signaling, lymphocyte and leukocyte activation, and wound healing. In contrast, myofibroblasts exhibited a less inflammatory profile, with significant upregulation in pathways related to ECM remodeling, leukocyte migration, wound healing, chemotaxis, and response to transforming growth factor beta (Supplemental Fig. 7F).

These findings indicate that High QASP expressing fibroblasts may drive a strong immune and inflammatory response, while High QASP myofibroblasts are more involved in wound healing and fibrosis.

## Discussion

In this study, we examined the transcriptional profiles of quiescent and senescent human cells to identify the QASP and distinguish it from the well-characterized SASP. Our scRNA-Seq and bulk RNA-Seq analyses and multiplex cytokine analysis challenge the traditional notion of quiescent as a dormant, non-inflammatory state, revealing instead that quiescent cells, particularly fibroblasts, express a distinct transcriptional profile marked by the unexpected upregulation of pro-inflammatory factors. This response varies by cell line, quiescence induction method, and oxygen levels. Moreover, we found that myofibroblasts in IPF tissues show increased expression of QASP genes, suggesting a potential role for quiescent cells in fibrotic processes. These findings position the QASP as an early inflammatory signature that may prime cells for senescence under chronic stress or aging conditions.

While quiescence is characterized by reduced metabolic activity and cell cycle arrest, our findings show that quiescent LF1 fibroblasts exhibit elevated expression of pro-inflammatory cytokines, including *IL6* and *IL1B*. This suggests that quiescent cells may contribute to tissue inflammation and degeneration even before entering senescence. Quiescent cells may adopt a pro-inflammatory state in response to environmental stresses, potentially influencing neighboring cells and initiating immune responses. These findings align with previous observations that quiescent cells play active roles in tissue maintenance and stress response [30].

Our transcriptomic comparative analysis across multiple human cell lines (BJ fibroblasts, IMR90 fibroblasts, LF1 fibroblasts and primary preadipocytes) and quiescence induction methods (CI or SW) reveals that the transcriptional profile of quiescent cells is both distinct from that of senescent cells and specific to the induction method used. These findings suggest that quiescence represents a heterogeneous state shaped by cellular context and external stimuli.

One of the most intriguing findings in our study was the substantial overlap between the QASP and the SASP, with 533 gene shared between the two gene sets. This overlap suggests that quiescent cells share functional characteristics with senescent cells, especially in the secretion of inflammatory and ECM remodeling factors. Mounting evidence shows that quiescent cells can transition directly into senescence, particularly under stress or with aging [12]. Our results support the idea that quiescence could act as a precursor or intermediate state to senescence. This view aligns with previous studies indicating that quiescence and senescence states are not entirely distinct and that cells in these states can be molecularly indistinguishable from each other at certain time points [31].

Our investigation into the effects of oxygen revealed its significant influence on QASP the expression in quiescent cells. Under physiological oxygen levels (2.5% O_2_), we observed a less pronounced QASP expression compared to atmospheric oxygen levels (20% O_2_), particularly in SW-induced quiescence. Cell cycle inhibitors like *CDKN1B* and *CDKN2C* were upregulated under low-oxygen conditions, suggesting that physiological oxygen may stabilize their expression. These findings imply that hyperoxia can enhance QASP expression, due to increased oxidative stress and ROS production. Given that tissue oxygen levels vary *in vivo*, the hyperoxic conditions typical in cell culture may artificially amplify inflammatory phenotypes [24, 32]. Our results underscore the importance of considering oxygen levels in cell culture studies, suggesting that physiological oxygen levels modulate quiescent cell function and influences their impact on the microenvironment.

Our cytokine analysis confirmed that the QASP is not merely a transcriptional signature but a functional secretory program. CI-induced quiescent cells secreted significantly higher levels of *IL6*, *IL1B*, *IL15*, and *CCL2* compared to both SW-induced quiescent and proliferating cells. This pattern mirrors the induction method-specific transcriptional regulation observed in QASP, reinforcing that quiescence is not a uniform state but rather a dynamic response shaped by external stimuli. Notably, *IL6* and *CCL2* secretion in CI exceeded the assay’s maximum detection limit, while proliferating cells were near the lower limit, precluding statistical significance despite substantial fold changes. This technical limitation likely underestimates the extent of cytokine secretion in CI-induced quiescence. Comparison of RNA-seq expression levels with cytokine secretion revealed that while transcriptional upregulation correlated with increased cytokine levels, although the relationship was not strictly proportional, suggesting additional post-transcriptional or post-translational regulation.

Finally, our analysis of scRNA-Seq data from IPF lung tissue revealed that myofibroblasts exhibit increased QASP gene expression compared to fibroblasts. This finding bridges our *in vitro* observations with physiological and pathological contexts, indicating that quiescent cells may contribute to fibrotic disease development or progression. Environmental stressors like mechanical strain, oxidative damage, or inflammatory signals in tissues could prompt quiescent lung fibroblasts to adopt a more pro-fibrotic, pro-inflammatory phenotype, contributing to the chronic inflammation associated with aging and playing a critical role age-related disease progression.

While our study offers valuable insights into the QASP and the progression from quiescence to senescence, several limitations should be noted. First, our cytokine analysis was restricted to a small panel, so extending our studies to a larger number of cytokines will further our understanding of the QASP. Second, the mechanisms underlying post translational QASP regulation in quiescent cells remain unclear, and further research is needed to elucidate the signaling pathways and regulatory networks involved.

Together, our results challenge the traditional view of cellular quiescence by highlighting its active role in modulating inflammatory responses to extrinsic stressors and contributing to tissue remodeling. The discovery of the QASP and its overlap with the SASP offers new insights into how quiescent cells may contribute to aging, tissue repair, and disease pathogenesis.

## Methods

### Cell Culture, Senescence and Quiescence Induction

#### For Single-cell RNA Sequencing

LF1 cells were isolated and established by the Sedivy lab and frozen samples were recovered and used.[33]. Proliferaring and senescent cell (RS, OIS, DDIS) single-cell data can be accessed from Gene Expression Omnibus database under accession number GSE173879 [16]. Proliferating cells were passaged in regular growth media and harvested at 60 – 80% confluency. Quiescent cells were obtained using a dual CI + SW method, by allowing cells to reach 100% confluency and subsequently incubating cells in serum free media for 2 days. RS cells were passaged in regular growth media until replicative exhaustion. These cells were maintained in the senescent state for 3–4 months. DDIS was induced by adding 40 µM etoposide (Selleckchem Cat#Catalog No.S1225) to LF1 regular media for 3 weeks, replacing the media every 3 days. OIS was induced in LF1 cells generated with a clonal cell line that possessed an ERT2-HRAS:G12V transgene. OIS was induced by adding 4-Hydroxytamoxifen (4-OHT) to regular growth media, at which point the cells underwent a hyperproliferative phase before becoming senescent after six days. Growth media consisted of HAMS F10, 15% FBS, and 1% Penicillin Streptomycin and Glutamine. Cells were maintained at 37°C at 5% CO_2_ and 2.5% O_2_. Cells were tested for mycoplasma contamination routinely using MycoAlert Mycoplasma Detection Kit (Lonza).

#### Bulk-RNA Sequencing

Male human diploid fibroblast cells (BJ), female diploid lung fibroblasts (IMR90) obtained from ATCC (Cat# CRL-2522 and CCL-186) and primary lung fibroblasts (LF1) obtained from the Sedivy lab were cultured. Primary human kidney preadipocyte cells were a gift from James Kirkland. BJ and IMR90 cells were grown in DMEM (Gibco Cat#11965) supplemented with 10%FBS and 1%penicillin streptomycin and glutamine. LF1 cells were cultured in Ham’s F10 (Gibco Cat#11550), 15% Fetal Bovine Serum (FBS), and 1% of penicillin streptomycin and glutamine. preadipocytes were grown in *α*MEM (Gibco Cat# 12571) and 10% FBS and 1%penicillin streptomycin and glutamine. For proliferating, cells were passaged in regular growth media and harvested at 60%–80% confluency. Quiescence was induced by contact inhibition by allowing the cells to reach full confluence then changing media every other day for 5 days, or by serum withdrawal by changing regular growth media to 0.1% FBS for 5 days. All cells were cultured in 37°C incubator with 92.5% N_2_ and 5%CO_2_. IMR90 cells were maintained at 2.5% O_2_ (low oxygen) or at 20%O_2_ (high oxygen). BJ cells were maintained at 20%O_2_. Preadipocytes were maintained at 20%O_2_. LF1 cells were maintained at 2.5% O_2_ (low oxygen). Cells were tested for mycoplasma contamination routinely using MycoAlert Mycoplasma Detection Kit (Lonza). Senescence induction on LF1 cells: LF1 cells frozen at a late passage were used to make replicative senescent cells.

Replicatively senescent LF1 cells were attained by serial passaging until cell division ceased. When cells reached 80% confluency, they were trypsinized and diluted 1:4. The time between passages for early cultures was around 4-5 days, however as cells approached senescence the duration between passages increased gradually to an interval of 2-3 weeks. After cells achieved 80% confluency, cells were trypsinized and diluted to 1:2 and this was designated as entry into senescence timepoint. 1 month after cells entered senescence, LF1 cells were harvested for the early replicative senescent cell state timepoint, and late replicative senescent cells were harvested 4 months after entry into senescence.

#### Senescence-Associated *β*-Galactosidase Staining

Cells were fixed and stained for senescence-associated *β*-galactosidase (SA-*β*-gal) activity using the classic X-gal protocol. Briefly, cells were washed in PBS, then fixed for 5 minutes in 2% formaldehyde and 0.2% glutaraldehyde in PBS. Following fixation, cells were incubated overnight at 37°C (no CO_2_) in staining solution containing 1 mg/mL X-gal, 5 mM potassium ferricyanide, 5 mM potassium ferrocyanide, 2 mM MgCl_2_, 150 mM NaCl, and 40 mM citric acid/sodium phosphate buffer (pH 6.0). Cells were washed and imaged the following day under brightfield microscopy.

### qPCR

Female human diploid lung fibroblast cells (IMR90) were used for this assay. Proliferating cells were passaged in regular growth media and harvested at 60 – 80% confluency. Contact inhibited quiescent cells were obtained by allowing cells to reach 100% confluency for 4 days in regular growth media. Serum withdrawal quiescent cells were obtained by incubating cells at 60% confluency in regular growth media for 0.01% FBS for 4 days. DNA Damage Induced Senescence (DDIS) was induced by adding 40 uM etoposide to LF1 regular media for 2 weeks, replacing the media every 3 days.Cells were maintained at 37°C at 5% CO_2_ and 2.5% O_2_ (low oxygen) or 20 O_2_ (high oxygen). RNA was extracted using the QIAGEN RNeasy micro kit. Single-stranded cDNA was synthesized from 1 µg of total RNA in a final volume of 20 µL using the QIAGEN QuantiTect Reverse Transcription Kit (Cat# 205311). cDNA was stored at 20,°C. A qPCR was performed on the cDNA using the Applied Biosciences Power SYBR Green PCR Master Mix and Applied Biosystems QuantStudio 6 Pro Real-Time PCR System machine according to manufacturer’s instructions. The primers used were from DeCecco et. al’s previously published study [34]. TBP was used as a control, as in Gonzalez et. al’s previously published study[35].

### Total RNA Generation

Total RNA was extracted from early passage, SW-induced quiescent, early and late senescent cells. Cells were washed with PBS and harvested using Trizol reagent (Invitrogen), after which Total RNA was harvested using the Qiagen Rneasy kit. NEBNext ultra II directional RNA library prep kit for Illumina along with the NEBNext Poly(A) mRNA magnetic bead isolation module was used to make isolated mRNA and cDNA libraries. Strand specific cDNA libraries were sent to Azenta for paired-end sequencing using Illumina HiSeq instrument. This experiment was performed using 4 biological replicates.

### RNA extraction and cDNA synthesis for Bulk RNA-Seq

Confluent BJ, IMR90, and preadipocyte cells were washed once with ice-cold PBS. Then, adherent cells were treated with 0.05% Trypsin and incubated at 37 °C for 3 minutes. Cells were then collected and centrifuged at 300 rcf for 3 minutes. RNA was collected following the Qiagen RNeasy Mini Kit (Cat#74104) protocol. The RNA concentration and purity were evaluated with a Qubit Broad Range Quantification Assay Kit (Q10211). RNA was submitted for Sequencing at Azenta (GeneWiz) with manufacturer recommended sequencing specifications.

### Bulk RNA-Seq Analysis

For the quiescent analysis, fastq files were received back from Azenta. For senescence analysis of BJ and IMR90 cells, fastq files were obtained from GEO Accession number GSE63577 [18]. Fastq files were assessed for quality using FastQC [36]. Reads were then aligned to reference genome hg38 using STAR. Subsequently, transcript quantification was performed using HTSeq [37] to obtain raw read counts per gene. PCA was conducted as a quality control measure to detect and exclude outlier samples. Specifically, for dataset quiescent IMR90, IHP2 was excluded. For dataset senescence high oxygen BJ and IMR90:, IM01 was excluded. For low oxygen LF1, Early Sen 4 was excluded (Supplemental Fig. 7). Batch effect correction was applied using the `removeBatchEffect` function from the `Limma package` [38]. Subsequently, raw counts were normalized and filtered using EdgeR. Differential expression analysis was performed using EdgeR’s generalized lineal model (GLM) approach [39]. Genes with FDR *<* 0.05 and log_2_FC *>*1 were considered differentially expressed. To analyze the expression of QASP genes, the differentially expressed genes were filtered using the predefined QASP list generated from the Secretome. Volcano plots, venn diagrams and heatmaps were generated using the `ggplot, ggVennDiagram` and `pheatmap` packages in R. GSEA [40] was performed to identify significantly enriched biological pathways in differentially expressed genes.

#### QASP UpSet Plot Construction

To visualize overlap in QASP gene expression, we used the `ComplexUpset` R package. Differential expression analysis was performed using the `limma` package on voom-transformed, batch-corrected RNA-seq data. Genes were filtered using an FDR *<* 0.05 and an absolute log_2_ fold change *>* 1. For each comparison (BJ_CI_, BJ_SW_, IMR90_CI_, IMR90_SW_, Preadipocyte_CI_, Preadipocyte_SW_, WI38_CI_), upregulated QASP genes were used to construct a binary presence/absence matrix across conditions. Intersections were visualized using `ComplexUpset` with horizontal bars and annotated intersection sizes. Only intersections containing *≥* 3 genes were displayed to reduce visual clutter.

### Single-cell RNA-Seq

LF1 cells were trypsinized and centrifuged at 500 rcf for 10 minutes. Cells were resuspended in cold phosphate-buffered saline and passed through a 40 uM Flowmi Cell Strainer. Cells were then counted and loaded onto the 10× chromium using V3 chemistry of 10× genomics’ 3-prime single-cell reagents with version 3 chemistry. Libraries were prepared according to the manufacturer protocol and sequenced on a NovaSeq platform at GeneWiz (Azenta) with manufacturer recommended sequencing specifications. Libraries were subjected to two rounds of sequencing.

### Single-cell RNA-Seq Analysis

FASTQ files were aligned to the hg38 human genome using CellRanger command-line tools, followed by secondary analysis with Seurat version 5 [41]. After filtering low-quality droplets, samples were processed in duplicate and merged post-filtering. Each cell population (Proliferating, quiescent, DDIS, OIS, RS) was filtered based on the number of genes, unique transcripts, mitochondrial gene expression, and GAPDH expression. Low-quality cells were clustered separately and excluded from further analysis. Filtered datasets were merged, scaled, and re-normalized using Seurat’s ScaleData, regressing out genes, transcripts, and mitochondrial content. Principal component analysis (PCA) was performed, with the top 20–30 components used for clustering via Seurat’s shared nearest neighbor Louvain algorithm. Differential expression was assessed using the MAST method in Seurat.

### Tissue Single-cell RNA-Seq Analysis

In this study, we utilized a lung single-cell RNA-seq dataset publicly available from Broad Institute Single-Cell Atlas [42]. GEO Series Accessions under GSE135893 [28]. Aligned reads were loaded to Seurat. The Seurat object was then filtered to keep only fibroblasts that were labeled “healthy” or “IPF” in the metadata. To distinguish between proliferating vs. quiescent vs. senescent fibroblasts, three filtering criteria were established. Proliferating cells were identified based on the expression of *MIK67, TOP2A*, or *PCNA*, with counts greater than zero being classified as proliferating. To identify senescent cells, a combination of criteria based on *CDKN2A* and *CKDN1A* expression levels was applied. Cells were identified as senescent if they met any of the following conditions:*CDKN2A >* 0, co-expression of *CDKN2A >* 0 and *CDKN1A >* 1, or *CDKN1A* expression *>* 4. Cells meeting any of these conditions were combined into a single set of senescent cells. Proliferating cells were excluded from the senescent category, and proliferating and senescent cells were excluded from the quiescent category. Violin Plots showing cell cycle expression markers and cell cycle inhibitors can be found in Supplemental Fig. 7A as well as the proportions of proliferating, quiescent and senescent fibroblasts and myofibroblasts in healthy vs. IPF tissue (Supplemental Fig. 7B).

To categorize the activation of fibroblasts into myofibroblasts, quiescent fibroblasts were isolated based on their Cell Cycle Category. A set of established myofibroblast-specific markers (*ACTA2, TAGLN, COL1A1, FN1, PDGFRB* and *POSTN* [43, 44] was defined. The expression of these markers was assessed using violin plots to establish a threshold, which was set at three. Cell expression any of the defined markers above the threshold were categorized as Myofibroblasts. To validate this classification, the expression levels of additional validation markers *CTHRC1, COL3A1, MMP9, ADAM12, CAV3, VIM, THY1* and *TGFB1* [45] was examined using violin plots.

High vs. Low QASP Module Score: To assess the relative expression QASP genes across cell populations, we developed a QASP module score using the Seurat R package (version 5.0.1). This score quantifies the average expression of our predefined set of QASP-related genes in single cells, normalized against a background of control genes to account for technical variability. Violing plots of the QASP score showed a bimodal distribution of Fibroblasts and Myofibloblasts. a threshold of 0.1 was chosen to categorize expression as “High QASP and Low QASP”. Differential gene expression was performed between these two groups for both cell lines, and significant (FDR *<* 0.05 and logFC *>*1) differentially expressed genes were extracted and used to perform GO term enrichment analysis.

### Multiplex Cytokine Assay

IMR90 cells were cultured in DMEM + 10% FBS except for SW, which were cultured in low serum media (DMEM + 0.1% FBS). Proliferating cells were passaged in regular growth media. Quiescent cell induction for CI was achieved by allowing the cells to reach full confluence fir 4 days, changing the media every other day. Serum withdrawal was achieved by allowing the cells to reach about 50% confluence and changing the media to low serum media for 4 days. Media was changed every other day. DDIS was induced by adding 40 µM etoposide (Selleckchem Catalog S1225) to IMR90 regular media for 2 weeks, replacing the media every 3 days. A day before collecting the conditioned media, all plates were rinsed with PBS and media was changed to serum free media for 24 hours for all conditions. Following 24 hours media was collected and filtered through a Millipore Steriflip Vacuum Tube Top Filter pore size 0.22 µm and stored at −80 C until the assay was performed

Cell culture medium samples were analyzed for inflammatory markers at the Fluid Biomarkers Laboratory of the Brown University Center for Alzheimer’s Disease Research. A custom Meso Scale Discovery (MSD) multiplex panel was designed using human U-PLEX Biomarker Group 1 markers (IFN-*γ*, IL-10, IL-15, IL-1*β*, IL-2, IL-4, IL-5, IL-6, MCP-1 (CCL2), and TNF-*α*) and was run according to the manufacturer’s instructions using a 2-fold sample dilution.

Briefly, each capture antibody was incubated with a U-PLEX-coupled linker for 30 min at room temperature (RT). Each reaction was stopped by incubation with Stop Solution for 30 min at RT. The linker-coupled capture antibodies were then combined, added to each well of the plates, and incubated at RT for 1 h with shaking at 700 rpm. Plates were washed three times with MSD Wash Buffer. Samples, standards, and controls were added and incubated at RT for 2 h with shaking. The plates were washed three times, followed by the addition of SULFO-TAG detection antibodies and incubation at RT for 1 h with shaking. Following three additional washes, MSD GOLD Read Buffer B was added to each well. Plates were read immediately on the MSD QuickPlex SQ 120MM Analyzer without further incubation.

Raw data were analyzed using MSD Discovery Workbench analysis software with a four-parameter logistic (4PL) model and 1*/y*^2^ weighting. Samples with a coefficient of variation greater than 20% were excluded from the analysis. Samples with concentrations below the assay’s limit of detection (LOD) were assigned a value equal to half the LOD. The LOD values (pg/mL) were as follows: IFN-*γ* (2.45), IL-10 (0.06), IL-15 (0.25), IL-1*β* (0.08), IL-2 (0.44), IL-4 (0.03), IL-5 (0.16), IL-6 (0.27), MCP-1 (0.22), and TNF-*α* (0.42).

Statistical analyses were performed using the edgeR R package. One-sided, one-sample *t*-tests were performed as appropriate based on the normality of the data. Statistical significance was defined as *p ≤* 0.05 and indicated by an asterisk (*).

## Supporting information

Supplementary Table S1. QASP and SASP gene lists and their reported overlap.

## Data Availability

The bulk RNA-seq and single-cell RNA-seq datasets newly generated for this study have been deposited in the NCBI Gene Expression Omnibus (GEO) under accession numbers GSE287058 and GSE287059, respectively. Publicly available datasets used in this study are identified in the Methods section.

## Author Contributions

A.R. and N.N. conceived and designed the study, S.E and A.R. prepared single-cell suspensions and prepared single-cell sequencing libraries for LF1 cells. A.R performed cell culture and RNA-extraction for bulk RNA-Seq for IMR90, BJ and preadipocytes. A.R performed analysis of publicly available bulk RNA-Seq data on senescent BJ and IMR90 cells. R.K. performed cell culture, RNA extraction and library preparation of LF1 bulk RNA-Seq data, S.E. and A.R. performed quality checks and filtering of scRNA-Seq data. A.R. performed quality checks, data integration, computational analyses and interpretation of all scRNA-Seq data, bulk RNA-Seq data and scRNA-Seq of tissue. A.R and N.N wrote and revised the manuscript.

## Competing Interests

The authors declare that they have no competing interests.

## Supplementary Figure Legends

**Figure S1.**
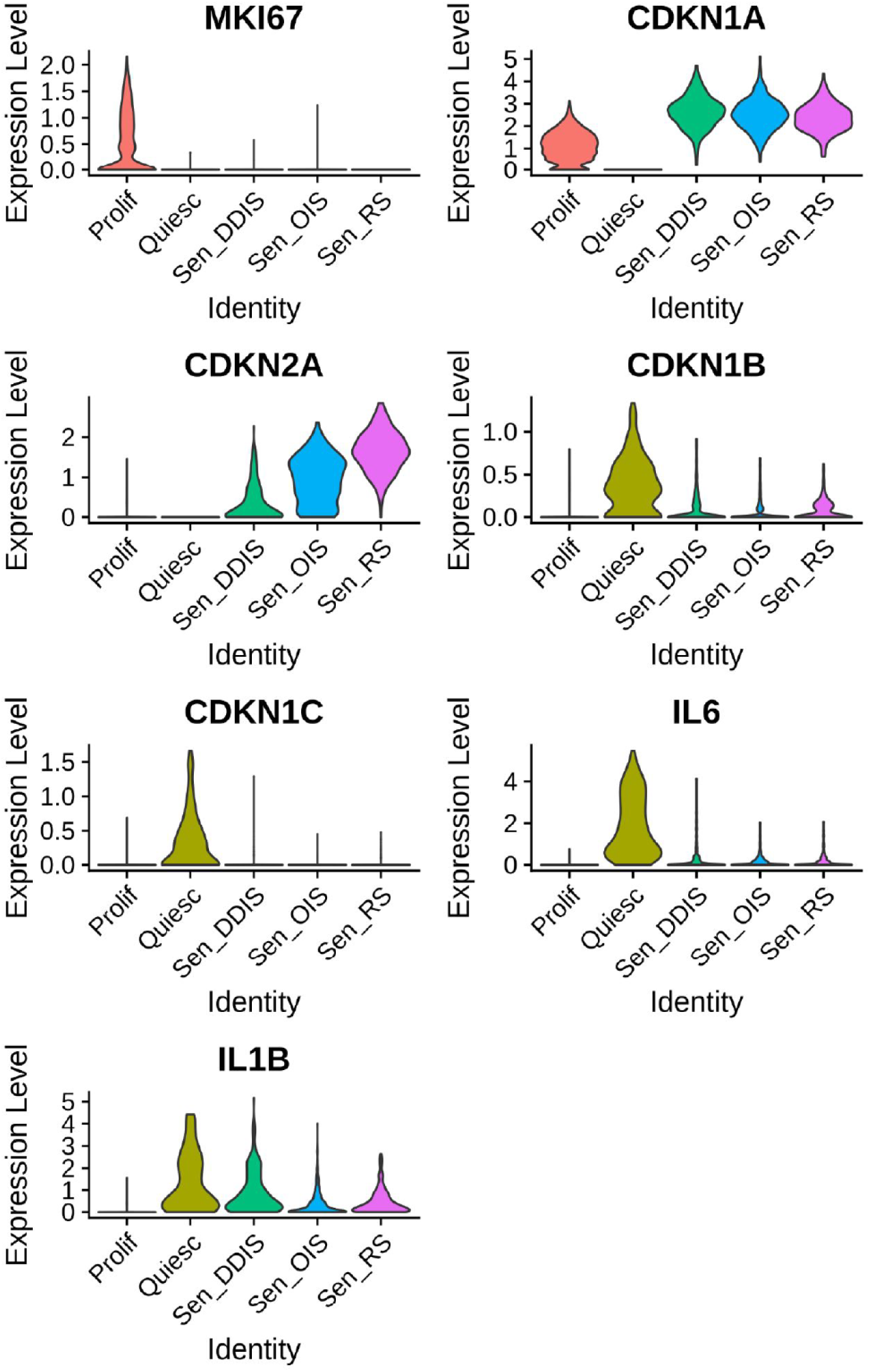
Single-cell analysis of quiescent and senescent samples. Violin plots showing expression levels of the proliferation marker *MKI67*, cell cycle arrest markers *CDKN1A*, *CDKN2A*, and *CDKN1B*, and inflammatory markers *IL6* and *IL1B* in *CDKN1A*- and *CDKN2A*-positive quiescent cells after filtering out *CDKN1A* and *CDKN2A*.

**Figure S2.**
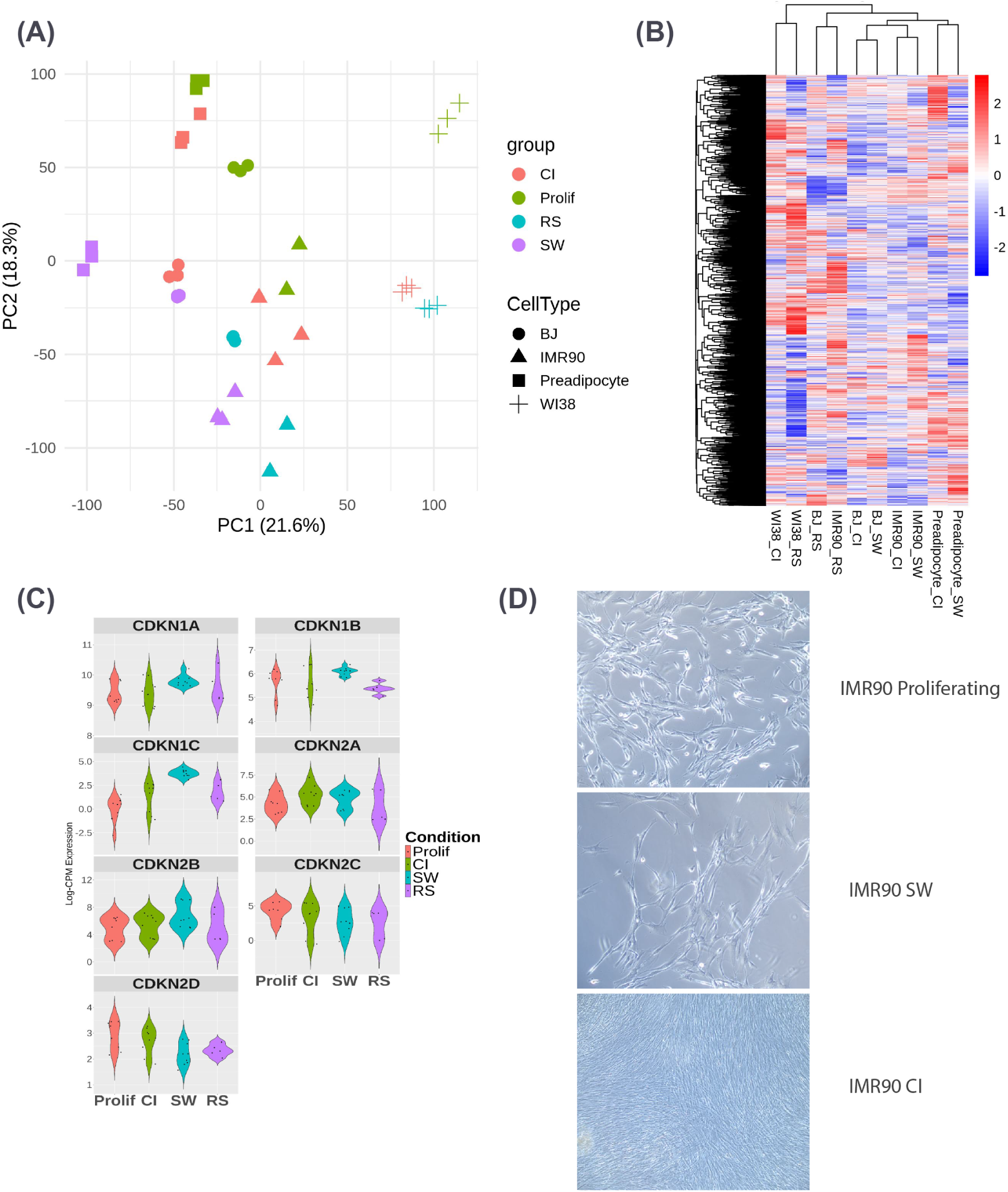
Continued analysis of high-oxygen BJ, IMR90, and preadipocyte samples. (A) PCA plot showing non-batch-corrected data for BJ, IMR90, and preadipocyte samples cultured under high-oxygen conditions, as well as senescent (RS) samples for BJ and IMR90 cells. Groups are distinguished by color and cell type by shape. (B) Heatmap displaying fold changes of differentially expressed genes (FDR *<* 0.05 and *|* log_2_ FC*| >* 1) across all contrasts for each cell line: CI vs. Prolif, SW vs. Prolif, and RS vs. Prolif (where applicable). (C) Violin plots depicting the upregulation and downregulation of cell cycle inhibitor genes (*CDKN2A*, *CDKN2B*, *CDKN2C*, *CDKN2D*, *CDKN1A*, *CDKN1B*, and *CDKN1C*) for each condition. (D) Senescence-associated *β*-galactosidase staining in IMR90 cells under proliferating, serum withdrawal (SW), and contact inhibition (CI) conditions. All samples were negative for SA-*β*-gal activity.

**Figure S3.**
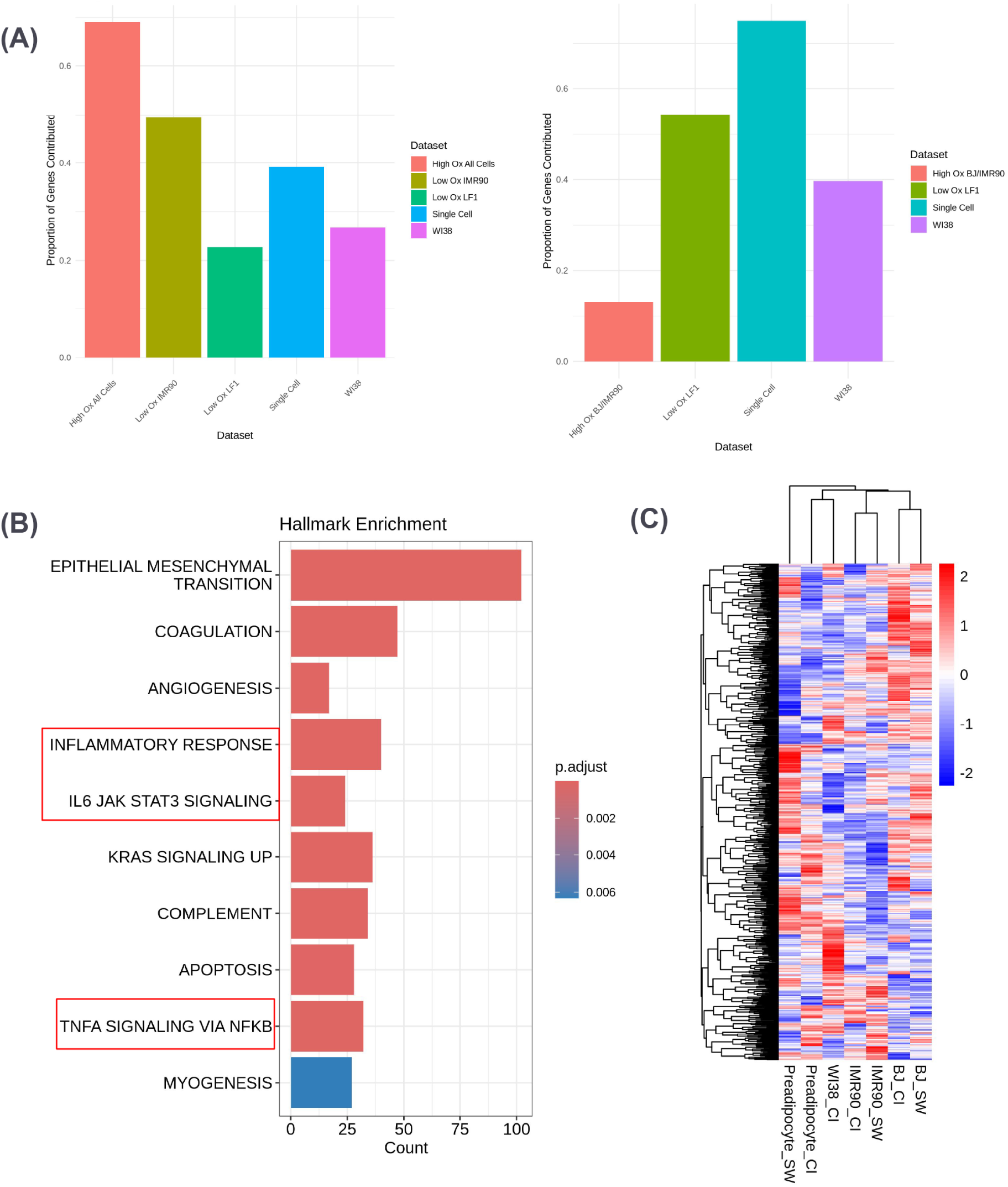
Characterization of the QASP and SASP. (A) Bar plots showing the proportional contribution of each dataset to the QASP (left) or SASP (right). Each condition is represented by a colored bar. QASP datasets include high-oxygen IMR90, BJ, and preadipocytes; low-oxygen IMR90; low-oxygen LF1; and single-cell LF1 quiescence datasets. SASP datasets include high-oxygen BJ and IMR90, low-oxygen LF1, and single-cell LF1 senescence datasets. (B) GO term enrichment analysis for the QASP gene set, revealing pathways associated with inflammatory responses, IL6-JAK-STAT3 signaling, and TNF-*α* signaling via NF-*κ*B. (C) Heatmap displaying differential gene expression (FDR *<* 0.05 and *|* log_2_ FC*| >* 1) of QASP genes across different cell lines and induction methods compared with proliferating cells.

**Figure S4.**
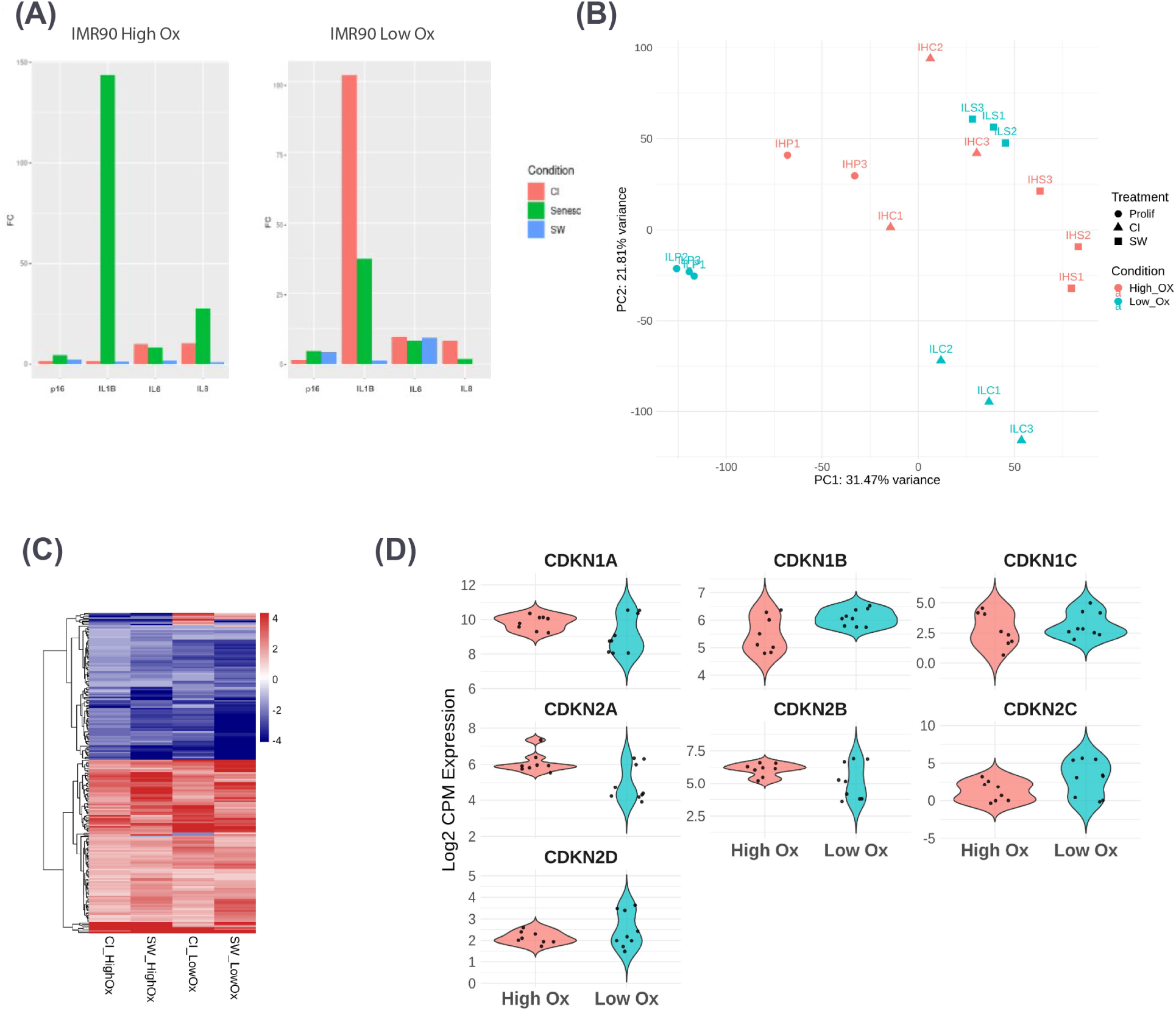
Effects of oxygen concentration on quiescence induction. (A) Bar plots of qPCR data showing gene expression changes under high-oxygen (20%, left) or low-oxygen (2.5%, right) conditions for *CDKN2A* (p16), *IL1B*, *IL6*, and *IL8* in quiescent cells (CI in red, SW in blue) or DDIS cells (green) for IMR90 cells. (B) PCA plot of bulk RNA-seq data showing sample clustering by condition and oxygen concentration for IMR90 cells. Low oxygen is depicted in blue and high oxygen in red. Proliferating cells are represented by circles, CI cells by triangles, and SW cells by squares. (C) Heatmap showing differentially expressed genes (DEGs) in quiescent IMR90 cells induced by CI or SW under low-or high-oxygen conditions. (D) Violin plots comparing the expression levels of cell cycle inhibitors (*CDKN2A*, *CDKN2B*, *CDKN2C*, *CDKN2D*, *CDKN1A*, *CDKN1B*, and *CDKN1C*) under high- and low-oxygen conditions.

**Figure S5.**
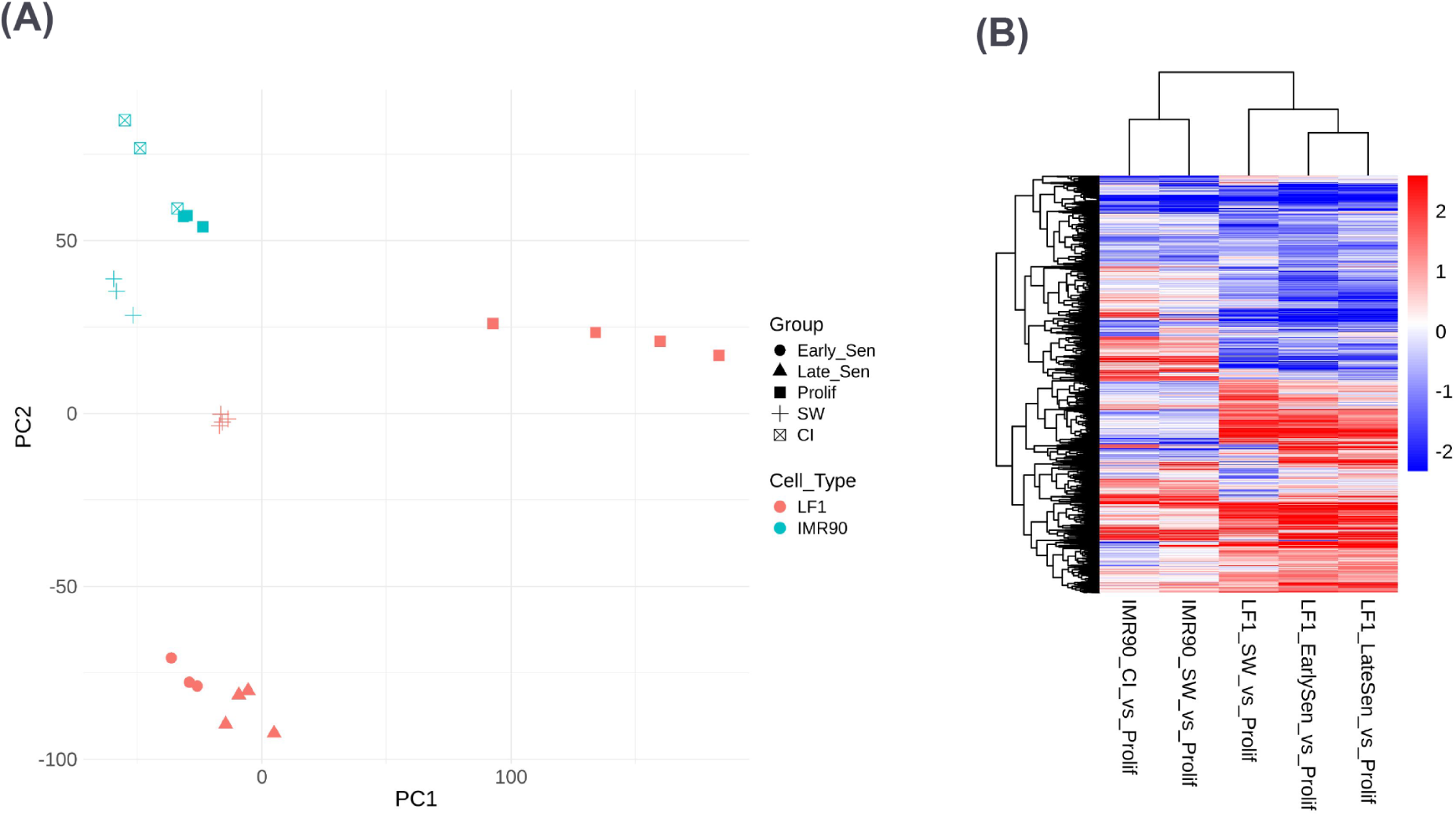
Non-batch-corrected PCA of low-oxygen IMR90 and LF1 cells. (A) PCA plot showing non-batch-corrected data from quiescent (CI or SW) IMR90 cells, SW LF1 cells, and senescent (early RS and late RS) LF1 cells cultured under low-oxygen (2.5%) conditions. Groups are distinguished by shape and cell type by color. (B) Heatmap showing significantly differentially expressed genes in quiescent IMR90 and LF1 cells or senescent LF1 cells compared with proliferating cells.

**Figure S6.**
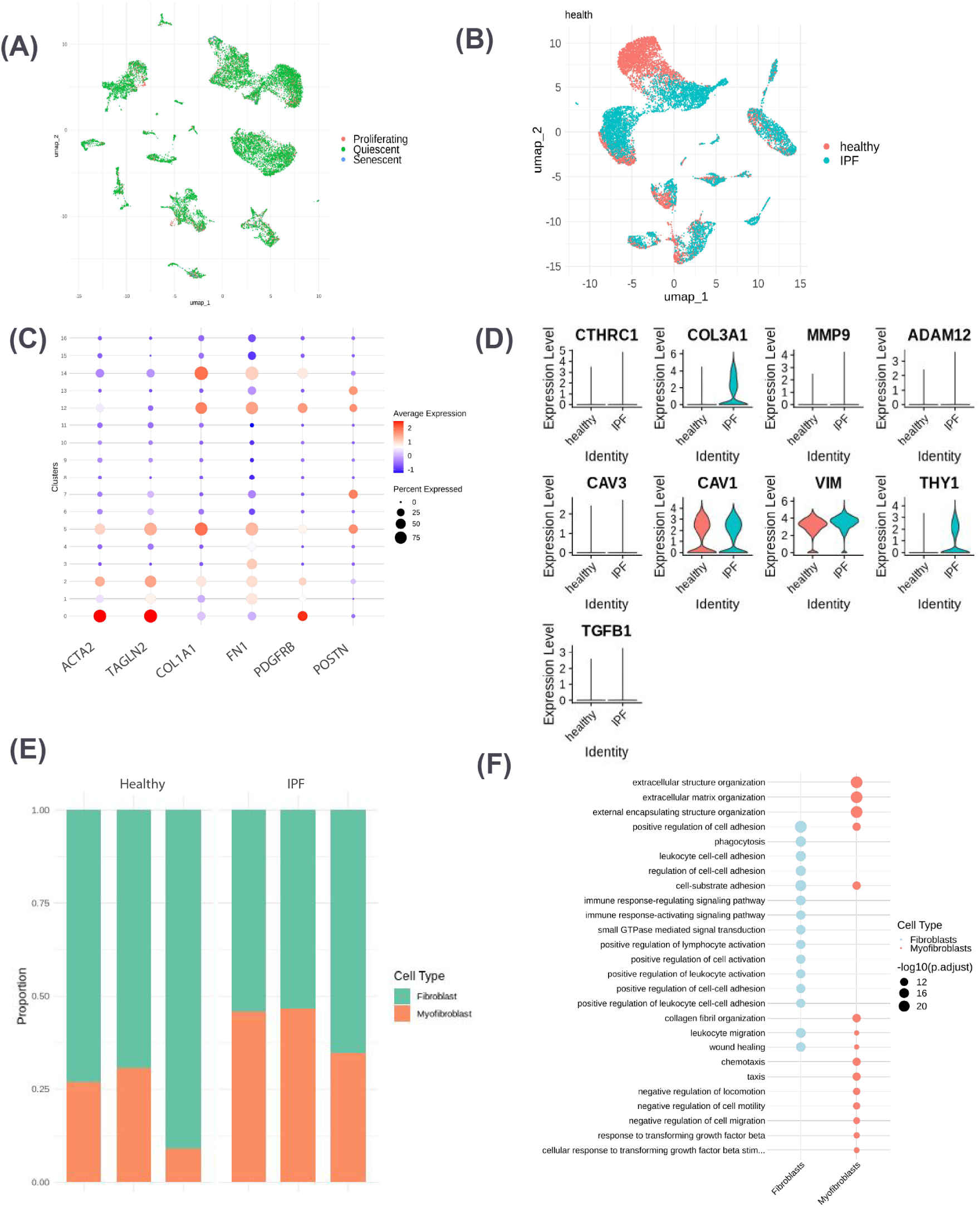
Analysis of quiescent fibroblasts and myofibroblasts in healthy and IPF lung samples. (A) UMAP of all fibroblasts from the single-cell RNA-seq dataset showing the distribution of cells classified as proliferating, quiescent, or senescent. Cell cycle categories were determined by specific marker expression. Proliferating cells are shown in red, quiescent cells in green, and senescent cells in blue. (B) UMAP of quiescent fibroblasts colored by health condition (healthy vs. IPF). Healthy cells are shown in red, while IPF cells are shown in blue. (C) Dot plot showing the expression of genes used to classify myofibroblasts across clusters (*ACTA2*, *TAGLN*, *COL1A1*, *FN1*, *PDGFRB*, and *POSTN*). Clusters 0, 2, 5, 12, and 14 were selected because they showed the highest expression levels. (D) Violin plots showing the upregulation of myofibroblast markers not used for myofibroblast classification (*CTHRC1*, *COL3A1*, *MMP9*, *ADAM12*, *CAV3*, *CAV1*, *VIM*, *THY1*, and *TGFB1*) in cells classified as myofibroblasts from healthy and IPF samples. (E) Bar plot comparing the proportions of myofibroblasts across cell cycle categories (proliferating, quiescent, and senescent) in healthy and IPF samples. Fibroblasts are depicted in green and myofibroblasts in orange. (F) GO term analysis showing pathways associated with differential expression between high-QASP and low-QASP populations in fibroblasts (blue) and myofibroblasts (red).

**Figure S7.**
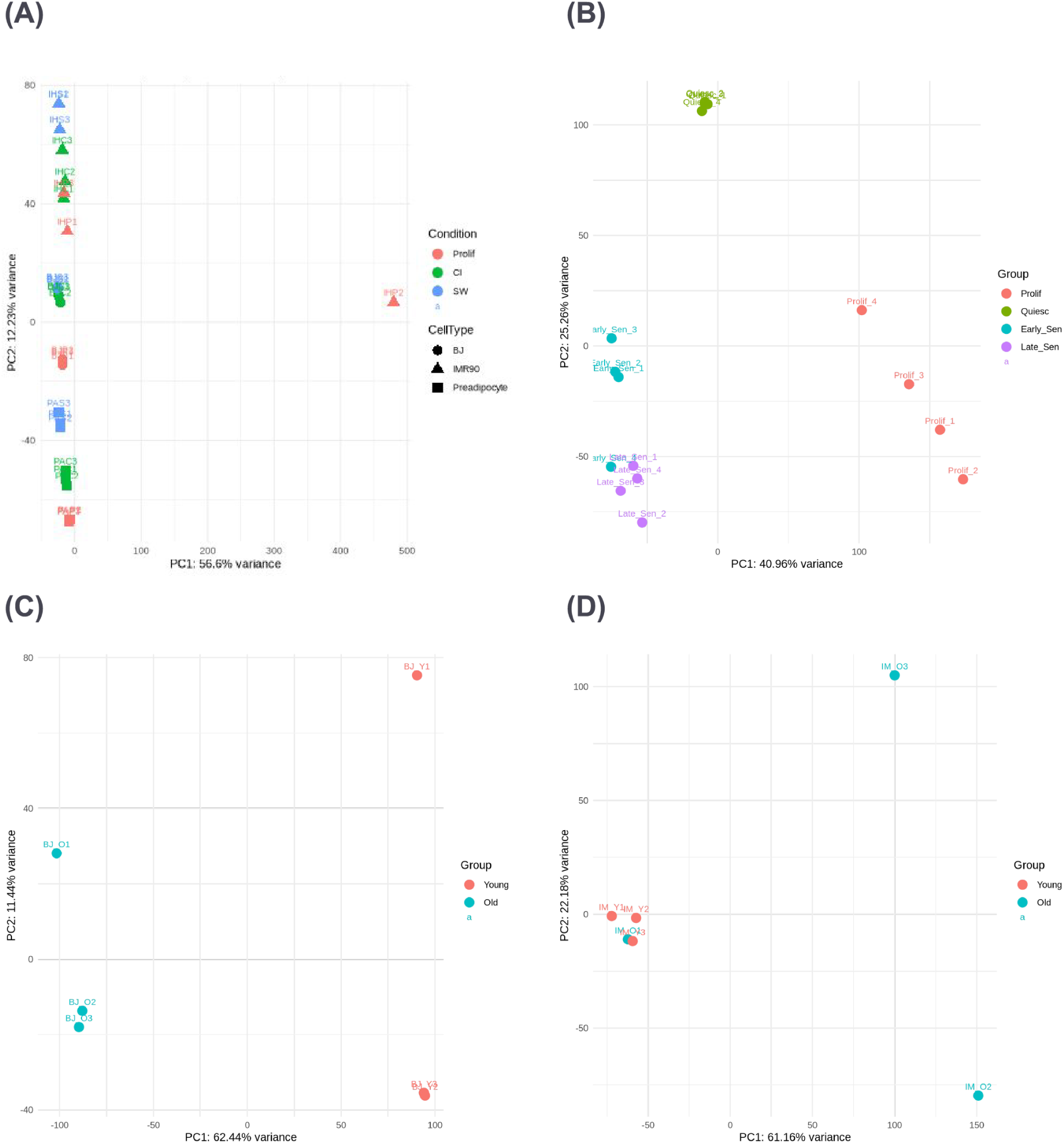
PCA filtering of outlier samples. (A) PCA plot of bulk RNA-seq data for quiescent BJ, IMR90, and preadipocyte samples, highlighting the outlier sample IHP2. This sample was excluded from further analyses. (B) PCA plot of quiescent and senescent LF1 cells, identifying Early Sen 4 as an outlier. Removal of this sample improved clustering and downstream analysis consistency. (C) PCA plot of senescent BJ cells. No significant outliers were detected, allowing the inclusion of all replicates in the analysis. (D) PCA plot of senescent IMR90 cells showing outlier clustering of sample IM01. This sample was excluded from further analyses.

## Supplementary Tables

**Supplementary Table S1. QASP and SASP gene lists and their reported overlap.**

The complete lists of SASP and QASP genes, together with the reported overlapping genes, are provided in Supplementary Table S1, available as a separate Excel file (Supplementary_Table_S1_QASP_SASP_Gene_Lists.xlsx).

## Notes

### Competing Interest Statement

The authors have declared no competing interest.

### Funding

- NIH: R21AG095984

